# Whole genome sequencing of pre-treatment and post-treatment locally advanced rectal cancer using long and short read technologies

**DOI:** 10.64898/2025.12.12.693949

**Authors:** Lauren McAuley, Lydia O’Sullivan, Liam Grogan, Adrian Murphy, Christina Skourou, Brendan Curran, Deborah McNamara, Joanna Fay, Brian O’Neill, Bryan T. Hennessy, Sinead Toomey, Simon J. Furney

**Affiliations:** Genomic Oncology Research Group, Department of Physiology and Medical Physics, RCSI University of Medicine and Health Sciences, Dublin, Ireland; St Luke’s Radiation Oncology Network, Dublin, Ireland; Department of Medical Oncology, Beaumont Hospital, Dublin, Ireland; Department of Surgery, RCSI University of Medicine and Health Sciences, Dublin, Ireland; Department of Pathology, RCSI University of Medicine and Health Sciences, Dublin, Ireland; Department of Medicine, RCSI University of Medicine and Health Sciences, Dublin, Ireland

**Author notes:** These authors contributed equally (Sinead Toomey, Simon J. Furney). Corresponding Author: Simon J Furney, Ph.D., Department of Physiology and Medical Physics, RCSI University of Medicine and Health Sciences, Dublin 2, Ireland.

## Abstract

We performed Illumina short-read and nanopore (ONT) whole genome sequencing (WGS) on samples obtained from a 37-year-old patient with locally advanced rectal cancer who did not respond to neoadjuvant chemoradiotherapy (nCRT). Pre-treatment and post-nCRT tumour biopsy, adjacent normal tissue, and matched pre-treatment blood samples were sequenced to identify somatic alterations. We used DNA methylation from ONT WGS to detect changes in the tumour epigenomes compared to adjacent normal tissue. We employed established pipelines for the identification of somatic alterations to assess the concordance of SNV, short indel, copy number and structural variation from the short-read and long-read tumour genome data.

Overall, 69.4% and 30.1% SNVs were concordant between technologies in the pre- and post-treatment tumours, respectively. A multinomial logistic regression was performed to further understand discordant SNV calls, highlighting tumour purity, presence in a repetitive region, strand bias, and germline variant allele frequency as factors contributing to discordance. Indel concordance was poor (26.8% and 9.2% in the pre- and post-treatment tumour, respectively). Copy number alteration profiles were highly similar while minimal concordance was reported for structural variants (2.4% and 0.3% for pre- and post-treatment tumours, respectively).

Finally, we used the ONT data to detect DNA modifications (5-methylcytosine and 5-hydroxymethylcytosine) between tumour and matched adjacent normal tissue. Based on genome-wide average CpG modification rates, global hypomethylation was observed in tumours compared to adjacent normal tissues. Biologically relevant epigenomic changes related to epithelial-mesenchymal transition were detected in the post-treatment tumour, as well as potential radiation-induced changes in the post-treatment adjacent normal tissue.

## INTRODUCTION

Genome sequencing has led to important revelations in the understanding and treatment of cancer. The first whole cancer genome sequence was published in 2008, where an acute myeloid leukaemia genome was generated using Illumina (ILMN) sequencing (Ley et al. 2008). It was realised that large-scale sequencing of cancer genomes would be necessary to capture the complexity of this disease, leading to the establishment of several genome sequencing initiatives such as the Pan-Cancer Analysis of Whole Genomes (PCAWG) Consortium, the Hartwig Medical Foundation, and the Cancer Programme of the 100,000 Genomes Project from Genomics England. These projects have generated large repositories of publicly available short read whole genome sequencing (WGS) data, including data from primary and metastatic tumours and matched normal tissues. These highly valuable resources have been used extensively by researchers, making substantial contributions to the current understanding of cancer genomics and assisting in the development of new computational tools to analyse short-read genome sequencing data (Aaltonen et al. 2020; Priestley et al. 2019; Martínez-Jiménez et al. 2023; Cameron et al. 2021; Sosinsky et al. 2024).

Short read sequencing using ILMN technology dominated the field of cancer genomics for two decades. However, inherent drawbacks of short reads include an inability to accurately align reads to repetitive regions and limited sensitivity when detecting structural variants (Guan and Sung 2016). In recent years, long-read sequencing technologies like nanopore sequencing (Oxford Nanopore Technologies; ONT) and HiFi sequencing (Pacific Biosciences; PacBio) have gained traction; long-read technologies typically produce reads over 10kb in length which enables the resolution of large repetitive regions and the characterisation of complex structural variants, effectively addressing the disadvantages of short-read sequencing. Recently, ILMN and ONT WGS have been used to characterise loss-translocation-amplification chromothripsis, a newly characterised mechanism of chromothripsis predominately occurring in osteosarcoma, with ONT WGS uncovering the allele-specificity of this mechanism (Espejo Valle-Inclan et al. 2025). ONT sequencing of a paediatric medulloblastoma resulted in the identification of templated insertions, a new class of structural variant (Rausch et al. 2023). An additional benefit of long read sequencing is the ability to directly obtain native epigenetic modifications from both DNA and RNA sequencing data without additional library preparation steps. While several techniques are available to detect DNA methylation from short reads, these involve harsh bisulfite treatments and/or enzymatic conversion steps (Liu et al. 2021; Wang et al. 2023; Füllgrabe et al. 2023). DNA methylation profiles generated from shallow nanopore WGS data have been successfully used for tumour classification (Vermeulen et al. 2023; Yuan et al. 2025). Moreover, long read RNA sequencing can sequence full length transcripts and detect RNA modifications such as m6A (Monzó et al. 2025).

Considering these advantages, as the accuracy of long-read technologies approach that of ILMN’s near-perfect reads and sequencing costs fall, the application of long-read sequencing to cancer genomics is becoming more viable. The Genomics England Cancer 2.0 and All of Us initiatives have launched pilot projects using ONT and PacBio sequencing, respectively, to assess how these technologies can improve the detection of small and large genomic variants and the feasibility of applying them in the clinical and research settings (Mahmoud et al. 2024). Several studies have utilised multiple sequencing platforms to generate call sets of structural variants in various cancer cell lines; for example, a truth set of structural variants was generated for the COLO829 melanoma cell line using Illumina, ONT, and PacBio sequencing platforms, with similar work from the SEQ2 Consortium generating a consensus set of structural variants in the HCC1395 breast cancer cell line (Espejo Valle-Inclan et al. 2022; Talsania et al. 2022; Nattestad et al. 2018; Aganezov et al. 2020). Additionally, the Genome in a Bottle Consortium has generated short and long read bulk WGS data of a matched tumour-normal pair from a pancreatic ductal adenocarcinoma to facilitate benchmarking of newly developed somatic variant callers (McDaniel et al. 2025). High quality call sets will be crucial for benchmarking new bioinformatic tools designed for use with long reads. Recent work utilising the Long-Read Personalised OncoGenomics (POG) cohort of 189 advanced cancer patients demonstrated the benefits of the addition of long read WGS data to the existing short read WGS and RNA-sequencing data available for this cohort; they showed that long reads enhanced complex SV detection and demonstrated how long-range phasing information can link allele-specific variants to gene expression (O’Neill et al. 2024). Long read sequencing technologies have great potential in the field of personalised medicine, however comparisons of the genomic variants detected by short and long read sequencing are needed to understand the limitations of available bioinformatic tools for long read data and enable accurate interpretation of this data (Liu et al. 2024).

Here, we present a pilot study that focused on comparing the abilities of these sequencing technologies to detect somatic variants in patient tumour genomes using available bioinformatic tools. To this end, ILMN and ONT WGS was performed on samples obtained from a patient with locally advanced rectal cancer (LARC) who did not respond to therapy. Pre-treatment and post-treatment tumour biopsies, pre-treatment and post-treatment adjacent normal tissues, and matched pre-treatment blood underwent WGS with both ILMN and ONT. Since adjacent normal tissue was available, DNA methylation from ONT WGS was also examined to detect changes in the epigenetic profile of the tumour and adjacent normal tissue.

## RESULTS

### Patient Characteristics

The patient was a 37-year-old male diagnosed with locally advanced rectal cancer. The patient underwent neoadjuvant chemoradiotherapy (nCRT) and completed 6 cycles of 5-fluorouracil and received 50.4Gy radiation over 28 fractions. The patient underwent surgical resection of the tumour 8 weeks after completing nCRT. The patient responded poorly to nCRT with a Mandard tumour regression grade of 4. Immunohistochemistry of mismatch repair proteins was performed with no loss of expression of PMS-2, MLH-1, MSH-2, or MSH-6, indicating that the tumour was microsatellite stable (MSS). Time to distal recurrence after beginning nCRT was 9 months. Metastases were observed in the liver and omentum. Time to local recurrence was 18 months after beginning nCRT. Overall survival was 23 months.

### Whole genome sequencing of tumour and adjacent normal tissues

WGS using short-read (ILMN) and long-read (ONT) sequencing technologies was performed on a total of 5 samples collected from one LARC patient, including pre-treatment and post-treatment primary tumour samples, pre-treatment and post-treatment adjacent normal tissues, and matched pre-treatment blood (Fig. 1A). Average coverage across all samples for ONT sequencing was 37X. For ILMN sequencing, average coverage across all samples was 45X (Supplementary Table 1). Whole genome sequencing data of tumour and adjacent normal tissues were compared to the matched blood sample to detect somatic single nucleotide variants (SNVs) and indels, somatic copy number alterations (sCNAs), and somatic structural variants (SVs). As adjacent normal tissues were available, changes in DNA methylation patterns between tumours and adjacent normal tissues were investigated.

**Figure 1.**
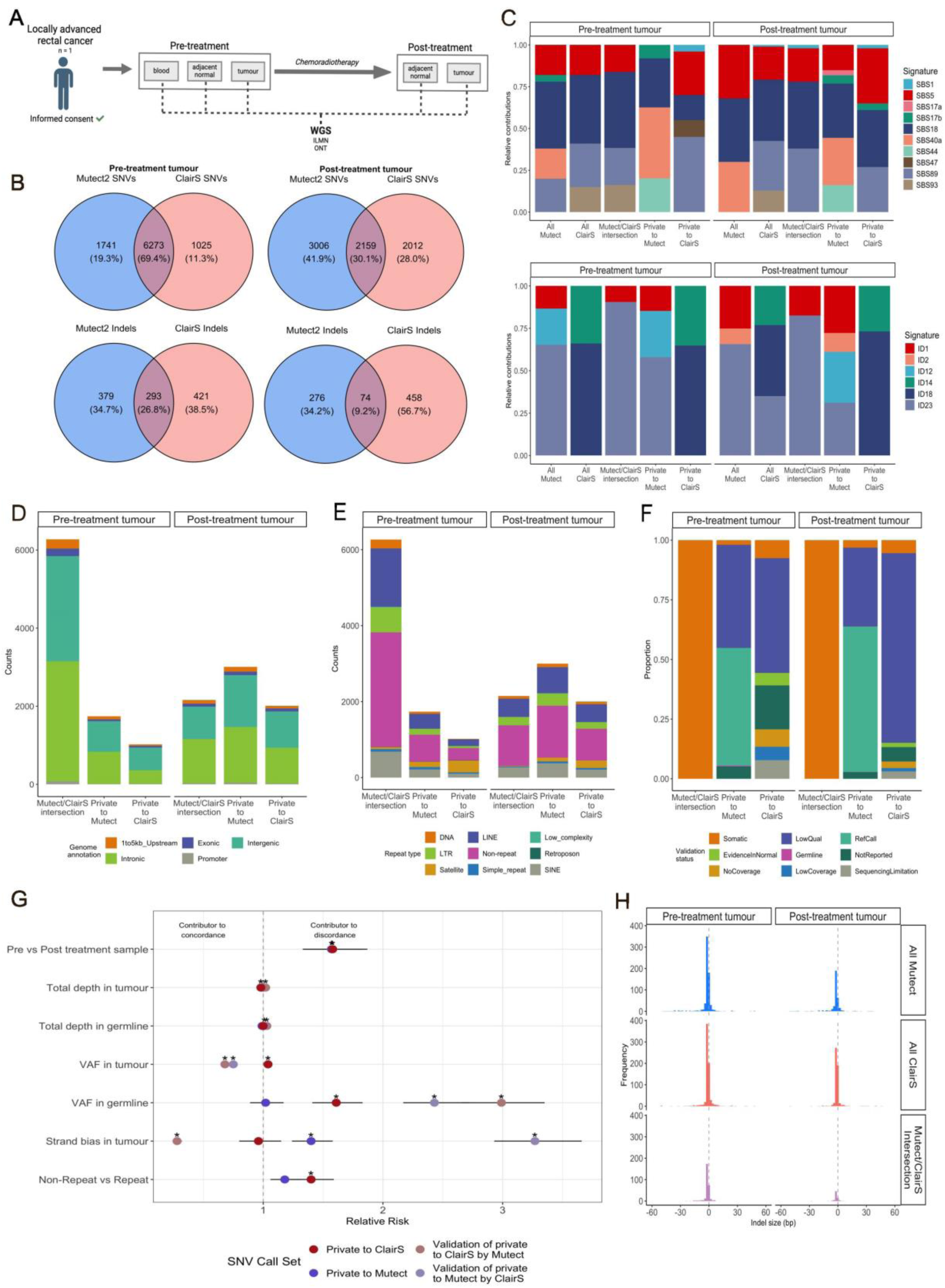
**A)** Schematic of the study design. **B)** Concordance of SNVs (top) and indels (bottom) between Mutect and ClairS for pre- and post-treatment tumours. **C)** SBS (top) and indel (bottom) signatures for each set in the pre- and post-treatment tumours. **D)** Genomic annotations of SNVs in each SNV set. **E)** Repetitive element annotation of SNV sets. **F)** Proportions of SNVs that were validated in orthogonal sequncing technologies. **G)** Multinomial regression analysis determined several factors, including VAF of the SNV in normal germline samples, contributed to discordance of SNV calls. **H)** Size distributions of indels identified by Mutect and ClairS.

### Concordance of small somatic variants between callers

#### SNVs

SNVs were detected by using Mutect in the ILMN data and ClairS in the ONT data (Table 1). The concordance of the SNVs detected between callers was examined for each sample (Fig. 1B), with 69.4% agreement in the pre-treatment tumour and 30.1% agreement in the post-treatment tumour. Few SNVs were concordant between callers in adjacent normal tissue samples (1.1% and 0.3% agreement in the pre-treatment and post-treatment adjacent normal tissues, respectively) (Supplementary Fig. 1A). The concordance of SNVs detected by a second short read small variant caller, Strelka2, was also assessed, with small differences in concordance observed with ClairS (Supplementary Fig. 1B). Mutect2 was chosen for subsequent analyses due to its better performance at variant frequencies below 10% (Chen et al. 2020). Sets of SNVs were defined as follows; all Mutect SNVs, all ClairS SNVs, SNVs detected by both Mutect and ClairS are termed ‘Mutect/ClairS intersection’ SNVs, while SNVs detected only in the ONT data or the ILMN data are referred to as ‘private to ClairS’ or ‘private to Mutect’, respectively.

**Table 1.** Counts of small somatic variants detected by Mutect and ClairS.

| Variant | Caller | Tumour |  | Adjacent normal |  |
| --- | --- | --- | --- | --- | --- |
|  |  | Pre-treatment | Post-treatment | Pre-treatment | Post-treatment |
| SNV | Mutect | 8014 | 5165 | 1018 | 450 |
|  | ClairS | 7298 | 4171 | 600 | 473 |
| Indel | Mutect | 672 | 350 | 124 | 117 |
|  | ClairS | 714 | 532 | 378 | 606 |

Mutational signature analysis of single base substitutions (SBS) was performed on the SNV sets. In both pre- and post-treatment tumours, SBS5 (clock-like) and SBS18 (damage by reactive oxygen species) were consistently detected across all Mutect, all ClairS and Mutect/ClairS intersection sets (Fig. 1C). SBS signatures detected in the set private to Mutect were largely comparable to the SBS signatures detected for all Mutect SNVs. A similar pattern was observed for the private to ClairS SNVs, however SBS18 was not detected in this set in the pre-treatment tumour.

The reasons behind SNV discordance between ILMN and ONT sequencing technologies were investigated using a total of 16,216 SNVs detected in tumour samples; 8,432 SNVs were in the Mutect/ClairS intersection set, 4,747 SNVs were private to Mutect and 3,037 SNVs were private to ClairS. To understand if genomic location affected SNV discordance, SNV sets were annotated with genomic regions as follows; 1-5kb upstream of a gene, exonic, intronic, intergenic, and promoter regions. A higher proportion of SNVs private to ClairS were in intergenic regions compared to other sets (1,508/3,037 (49.7%) private to ClairS, 2,113/4,747 (44.5%) private to Mutect, and 3,535/8,432 (41.9%) in the Mutect/ClairS intersection) (Fig. 1D). As repetitive elements are known to pose difficulties for variant detection, SNV sets were annotated with various repetitive elements to assess differences in the abilities of the sequencing technologies to profile these regions (Fig. 1E). SNVs in satellite regions were more prevalent in the set private to ClairS (503/3,037 (16.6%) in private to ClairS set vs 232/4,747 (4.9%) private to Mutect set vs 58/8,432 (0.7%) in the Mutect/ClairS intersection set).

A multinomial logistic regression was performed to further understand the factors contributing to discordant SNV calls in tumour samples. Evidence of SNVs that were private to a caller (i.e. private to a sequencing technology) was extracted from the orthogonal data type where a call was not made (termed validation calls). Total read depths, variant allele frequencies, and strand bias information was obtained for a given SNV in both ONT and ILMN data. This data was available for 13,831/16,216 SNVs; 8,432 SNVs were in the Mutect/ClairS intersection, 2,599 SNVs private to ClairS were validated in ILMN data and 4,574 SNVs private to Mutect were validated in ONT data (Fig. 1F, Supplementary Table 2). This variant information, along with sample and repetitive region annotations, was used as input for the regression model. Several factors contributed to SNV discordance between callers/sequencing technologies (Fig. 1G). Sample was used as a surrogate for tumour purity and was a contributor to discordance for both private SNV sets (SNVs private to Mutect RR = 1.57, p < 0.001; SNVs private to ClairS RR = 1.58, p <0.001, which was in line with the disparity in concordance initially observed. Similar proportions of SNVs were in repetitive regions across the SNV sets (1,542/2,599 (59.3%) private to ClairS, 2,527/4,574 (55.2%) private to Mutect, and 4,348/8,432 (51.6%) in Mutect/ClairS intersection). Whether an SNV was in a repetitive region contributed to discordance, with SNVs private to ClairS 40% more likely to be discordant for this reason (RR = 1.40, p < 0.001). This was not a significant factor for discordance in the SNVs private to Mutect (RR = 1.18, p = 0.10). Additionally, strand bias in the ONT validation data of SNVs private to Mutect contributed to discordance of these SNVs (ONT validation of private Mutect SNVs; mean strand bias = 0.75, RR = 3.27, p < 0.001, private Mutect SNVs; mean strand bias = 0.57, RR = 1.40, p < 0.001). Germline support for alternate alleles also contributed to SNV discordance; ClairS calls in the ONT data were 61% more likely to be discordant based on VAF in the germline sample (RR=1.61, p< 0.001), with validation of these calls in ILMN data showing an increased likelihood of discordance when considering VAF in the germline sample (RR = 2.99, p < 0.001). Similarly, validation of Mutect SNVs in the ONT data showed an increased likelihood of discordance due to VAF in the germline sample (RR = 2.43, p < 0.001).

#### Indels

Concordance of indels between Mutect and ClairS was 26.8% (293/1,093) in the pre-treatment tumour and 9.2% (74/808) in the post-treatment tumour (Fig. 1C). Concordance was minimal between callers in adjacent normal tissues (Supplementary Fig. 2A). Interestingly, most concordant indels in tumours were of the same length and position (Fig. 1H). Of all indels detected in the pre-treatment tumour, 428/672 (63.7%) and 451/714 (63.2%) detected by Mutect and ClairS, respectively, overlapped (minimum 1bp) or were within 5bp of a homopolymer. In the post-treatment tumour (195/350 (55.7%) of all Mutect indels and 368/532 (69.2%) of all ClairS indels overlapped or were within 5bp of a homopolymer) (Supplementary Table 3). Indel concordance in the pre- and post-treatment tumours increased to 40.8% and 15.6%, respectively, when indels overlapping/adjacent to homopolymers were excluded (Supplementary Fig. 2B). Of concordant indels detected between technologies in the pre- and post-treatment tumours, 146/293 (49.8%) and 31/74 (41.2%) overlapped or were adjacent to homopolymers indicating that homopolymers were not a barrier to accurate indel detection in this case.

Similarly to SNV sets, indel sets were defined. Indel signatures were extracted using all indels detected as input regardless of homopolymer status (Fig. 1C). ID18 (colibactin exposure) and ID14 (aetiology unknown) were consistently detected by ClairS across tumours, while ID1 (slippage during DNA replication) and ID23 (aristolochic acid exposure) were consistently detected by Mutect. ID12 (unknown aetiology) was also detected in tumours by Mutect, along with ID2 (slippage during DNA replication) in the post-treatment tumour. Mutect/ClairS intersection signatures more closely resembled signatures extracted from Mutect indels as ID1 and ID23 were detected in this indel set.

### Concordance of small somatic variants between tumours

Sequencing of tumours with short and long read technologies enabled a comparison between technologies to detect tumour evolution. Concordance of SNVs between tumours was similar between technologies with 43.5% and 41.1% of SNVs detected in both the pre- and post-treatment tumours by Mutect and ClairS, respectively (Fig. 2A). Of all 10,666 SNVs detected across the tumour samples by both Mutect and ClairS, 1,932 (18.1%) were common to pre- and post-treatment tumours sequenced by ILMN and ONT (Fig. 2B). A total of 2,920 SNVs were detected by both sequencing technologies in a given tumour but detected by a single sequencing technology in the other tumour, i.e. detected in 3 of 4 samples. In some of these cases, low quality evidence from an orthogonal sequencing technology supported these SNVs in the remaining tumour; including these SNVs increased concordance to 48.3% and 28.6% in the pre- and post-treatment tumours respectively, with an increase in overall concordance to 28.0% (2,988/10,666) (Supplementary Fig. 3A, B).

**Figure 2.**
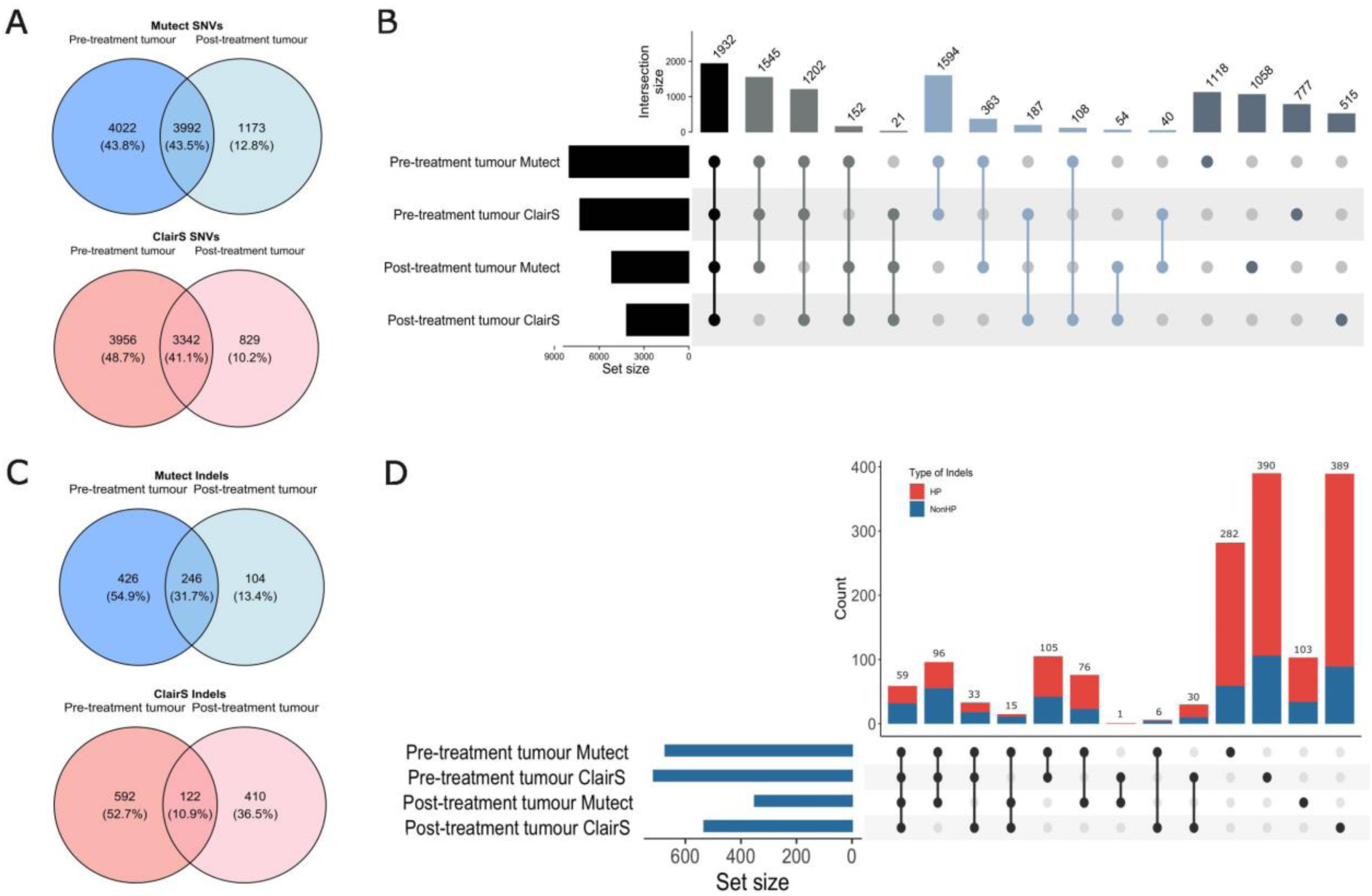
**A)** Concordance of SNVs detected by Mutect and ClairS in pre-treatment and post-treatment tumours. **B)** UpSet plot representing concordance of SNVs detected by Mutect and ClairS across pre-and post-treatment tumours. **C)** Concordance of indels detected by Mutect and ClairS in pre-treatment and post-treatment tumours. **D)** UpSet plot representing concordance of indels detected by Mutect and ClairS across pre- and post-treatment tumours, with indels overlapping or adjacent to homopolymers (HP) in red and indels not overlapping/adjacent to homopolymers (Non-HP) in blue.

Indel concordance between tumours was measured across all indels and when excluding indels overlapping or adjacent to homopolymers. Of all indels detected in the pre- and post-treatment tumours, 59/1,585 (3.7%) indels were detected in both the pre- and post-treatment tumours by both Mutect and ClairS (32/483 (6.6%) when excluding indels overlapping/adjacent to homopolymers) (Fig. 2C). A further 144/1,585 (9.1%) of indels were detected in the pre- and post-treatment tumours but were not detected by one orthogonal technology in one tumour; 84/483 (17.4%) were concordant when excluding indels overlapping/adjacent to homopolymers (Fig. 2D). A total of 105 indels (6.6%) were detected by Mutect and ClairS in only the pre-treatment tumour (42/483 (8.7%) when excluding indels overlapping/adjacent to homopolymers), while no indels private to the post-treatment tumour were validated by an orthogonal sequencing technology.

### Somatic Copy Number Alterations

Somatic copy number alterations (sCNAs) were detected using FACETs on the ILMN data and Wakhan on the ONT data. Ploidy estimates between orthogonal sequencing data were highly similar for the pre-treatment tumour with estimates of 2.56 and 2.53 for ILMN and ONT, respectively, with similar ploidy estimate in the post-treatment tumour (2.45 and 2.49 for ILMN and ONT, respectively). Additionally, purity estimates based on ONT data were notably higher compared to the ILMN data (0.63 and 0.54 for ONT vs 0.4 and 0.21 for ILMN for the pre- and post-treatment tumours, respectively).

Similar sCNA patterns were evident between ILMN and ONT WGS data (Fig. 3A). Hyperploidy of multiple chromosomes (1q, 2, 6, 7, 8, 12, 13, 14, 16, 19, 20) was consistently reported across technologies and tumours. A genome is classified as having undergone WGD if over 50% of the autosomal genome has a major copy number of at least 2 (Bielski et al. 2018). Using this metric, it was determined that the pre-treatment tumour had undergone a WGD event. Copy number signature analysis also supported a past WGD event; CN1 (diploid) and CN2 (tetraploid) signatures were detected in the FACETS and Wakhan segmentation data of pre- and post-treatment tumours (Fig. 3B). Co-occurrence of these signatures indicates a sub-tetraploid copy number profile (Steele et al. 2022). Contributions of CN2 were lower in Wakhan and CN23 (an oversegmentation artefact) was detected (Fig. 3C). CN23 is a result of segments <100kb and is typically observed in CN data generated by SNP6 arrays. Segment profiles derived from Wakhan displayed an increased number of segments <100kb compared to the FACETS segments profiles, though whether this is a sequencing artifact or genuine CN difference remains unclear (Fig 3C). Consistent with histopathological findings of microsatellite stability in this tumour, the CN signature linked to deficient mismatch repair (CN25) was not detected in short or long read WGS data.

**Figure 3.**
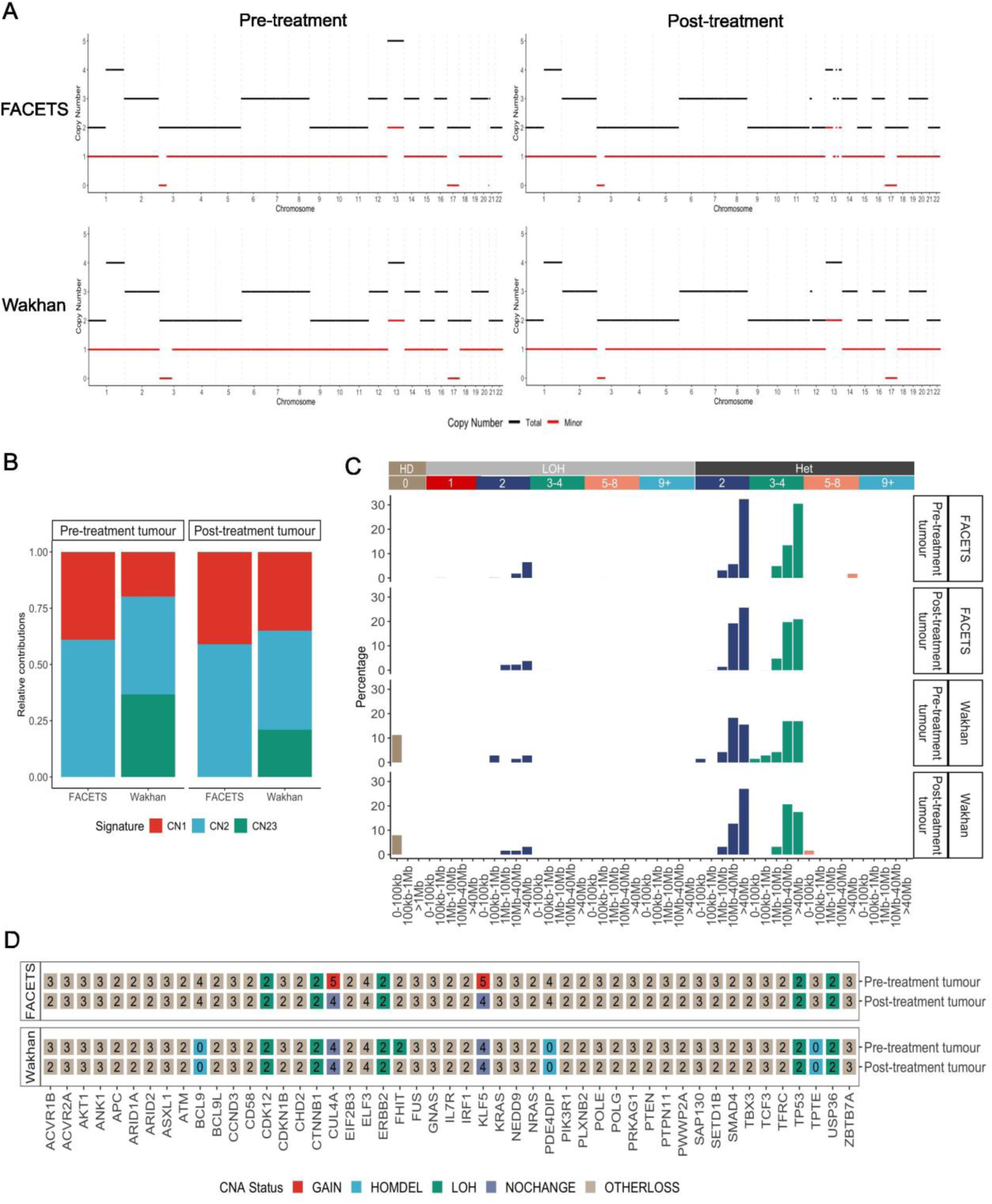
**A)** FACETS and Wakhan copy number profiles for the pre- and post-treatment tumours **B)** Copy number signatures detected in the pre- and post-treatment tumours using segmentation profiles from FACETS and Wakhan. **C)** Mutational profile of copy number signatures detected in pre- and post-treatment tumours by FACETS and Wakhan. **D)** Copy number status of driver genes.

Copy number changes were defined based on the occurrence of WGD as described previously (Cornish et al. 2024). The timing of sCNAs can be deduced from the total and minor copy numbers. FACETs and Wakhan detected LOH of the entirety of chromosome 17 and of a 41.56Mb region of chromosome 3 (chr3p26.3-p22.1) in the pre-treatment tumour, which occurred prior to WGD. This patient was 37-years old; the observation of LOH of TP53 prior to a WGD event is consistent with previous findings linking TP53 loss and WGD to early onset CRC (Kim et al. 2021). Additionally, a loss occurred in chromosome 1q along with a gain in chromosome 13 occurred in the pre-treatment tumour after WGD. Subsequent losses of single copies of chromosomes 2, 6-8, 12, 14, 16, 19, 20 and double losses in chromosomes 1p, 3-5, 9-11, 15, 17, 18, 21 and 22 were observed. Copy number states in the post-treatment tumour remained predominately stable compared to the pre-treatment tumour, however the majority of chromosome 12 experienced a loss with the exception of a 15.64Mb region containing *KRAS*. Additionally, single and double losses of segments of chromosome 13 are observed in the post-treatment tumour sequenced by ILMN; however, this pattern is not evident in the ONT data, potentially as Wakhan did not detect any subclonal sCNA events.

Copy number alterations were observed in several driver genes previously reported to experience copy number changes in primary MSS stable CRC and in driver genes in the Cancer Gene Census (Fig. 3D) (Cornish et al. 2024; Sondka et al. 2018). Loss of heterozygosity in *CDK12*, *CTNNB1*, *ERBB2*, *TP53*, and *USP36* was consistently detected by short and long read technologies, while gains were less consistently identified. Homozygous deletions of *BCL9*, *PDE4DIP*, and *TPTE* were observed in long reads but not in short reads; however, upon manual inspection coverage across these genes is high and these calls appear to be false positives homozygous deletions. Potentially, phasing in these regions was not optimal and may have contributed to false positive calls.

### Structural Variants

While short reads can reliably detect most large (>10kb) somatic structural variants within mappable regions, long reads are better able to detect smaller structural variants, insertions, and complex SVs due to their improved mappability and haplotype-specificity (Choo et al. 2023; Keskus et al. 2025). Here, somatic SVs were detected using Manta and Severus in ILMN and ONT data, respectively. In line with previous observations that long reads improve the ability to detect smaller SVs, we report that an increased number of smaller SVs (< 10kb) was detected by Severus compared to Manta (Choo et al. 2023). Additionally, Severus detected a markedly greater number of insertions and this observation was consistent across all tumour samples (Fig. 4A, Supplementary Fig. 4A). Additionally, few large somatic SVs were detected which is consistent with previous work (Zeng et al. 2025). We note that the proportion of classes of SVs detected in ILMN and ONT data differed; while just a single insertion was detected across both ILMN tumour samples, this SV class was the most prominent class detected in ONT samples (Table 2). The majority of SVs detected by Manta across ILMN tumour samples were breakends (BNDs), many of which can be attributed to translocations (Fig 4B). It is important to note that Severus reports translocations as insertions.

**Figure 4.**
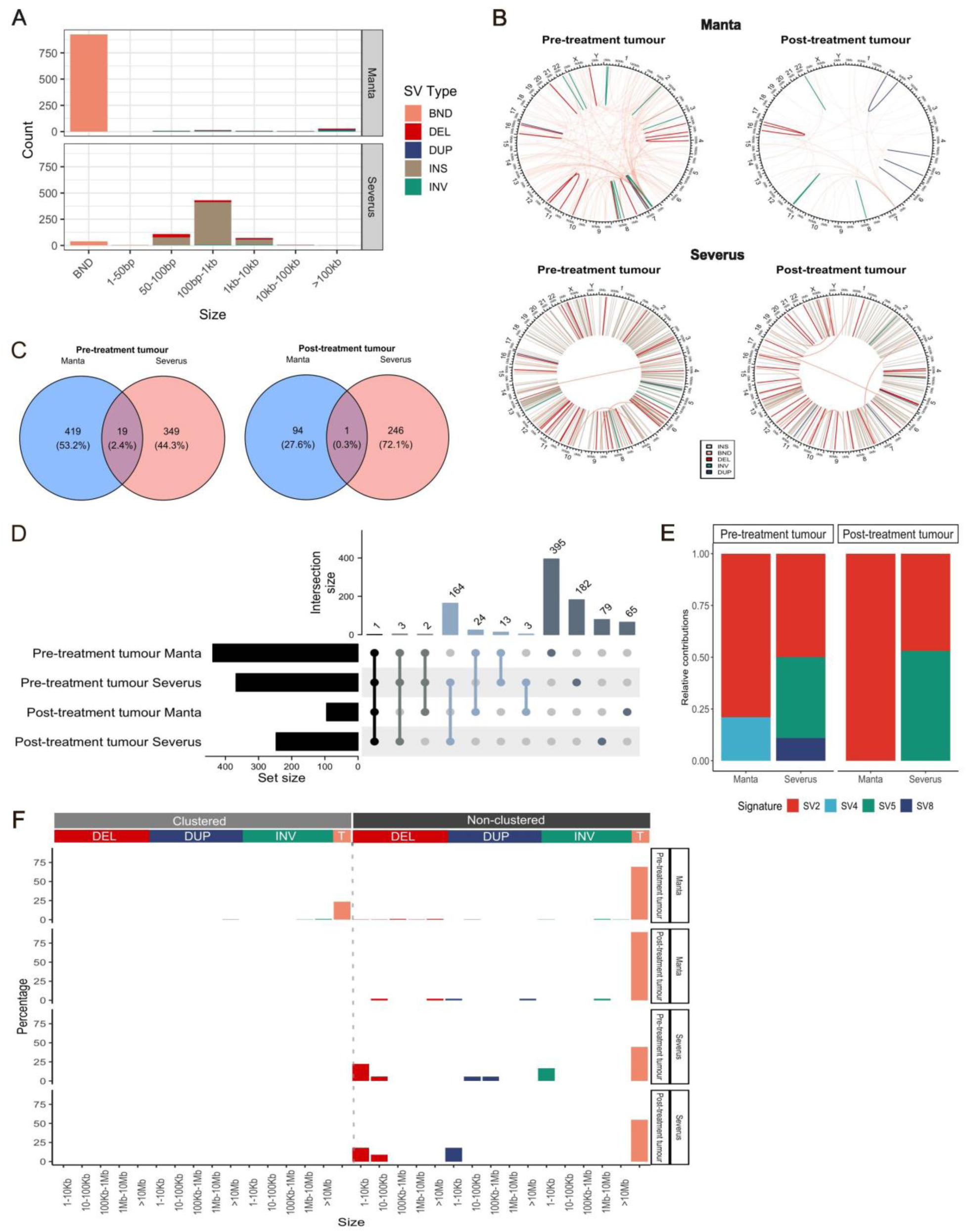
**A)** Distribution of SV sizes detected across all samples by Manta and Severus. **B)** Structural variants detected in the pre- and post-treatment tumours by Manta and Severus. **C)** Minda concordance of SVs between Manta and Severus for the pre- and post-treatment tumours. **D)** Minda concordance of SVs across all tumours and SV callers. **E)** Relative contributions of SV signatures differed between tumour samples snd SV callers. **F)** Mutational profile of SV signatures.

**Table 2.** Counts of structural variants detected by Manta and Severus.

| Caller | SV class | Tumour |  | Adjacent normal |  |
| --- | --- | --- | --- | --- | --- |
|  |  | Pre-treatment | Post-treatment | Pre-treatment | Post-treatment |
| **Manta** | **Total** | **438 (100)** | **95 (100)** | **324 (100)** | **129 (100)** |
|  | INS | 1 (0.2) | 0 (0) | 0 (0) | 0 (0) |
|  | DEL | 13 (3.0) | 2 (2.1) | 1 (0.3) | 4 (3.1) |
|  | BND | 412 (94.1) | 86 (90.5) | 316 (97.5) | 112 (86.8) |
|  | INV | 10 (2.3) | 3 (3.2) | 4 (1.2) | 7 (5.4) |
|  | DUP | 2 (0.5) | 4 (4.2) | 3 (0.9) | 6 (4.7) |
| **Severus** | **Total** | **368 (100)** | **247 (100)** | **35 (100)** | **9 (100)** |
|  | INS | 313 (85.1) | 199 (80.6) | 14 (40.0) | 4 (44.4) |
|  | DEL | 27 (7.3) | 27 (10.9) | 9 (25.7) | 2 (22.2) |
|  | BND | 16 (4.3) | 7 (2.8) | 5 (14.3) | 0 (0) |
|  | INV | 10 (2.7) | 2 (0.8) | 1 (2.9) | 1 (11.1) |
|  | DUP | 2 (0.5) | 3 (1.2) | 0 (0.0) | 1 (11.1) |

Comparison of structural variants is not straightforward due to differences in how variant calling tools denote these variants. An SV benchmarking tool, Minda, was used to compare SVs between Manta and Severus. When comparing two SVs, Minda will declare them as the same variant if both the start and end positions of each variant are within a given window between samples (Keskus et al. 2025). Concordance rates between technologies were low; only 19 (2.4%) SVs detected by Manta and Severus in the pre-treatment tumour were concordant using Minda (Figure 4C). Upon visual examination of circos plots, it was evident that concordance was likely higher. For example, Manta detected a high density of translocations between chromosome 13 and chromosome 7 in the pre-treatment tumour (Fig. 4B). In the same region of chromosome 13, Severus detected a similar hotspot of insertions in the pre-and post-treatment tumours. Comparison of genomic coordinates found that for 7 of 24 insertions detected by Severus in chromosome 13 of the pre-treatment tumour, Manta detected breakends within 500bp of the insertion start position (4/7 had breakends within 10bp of the insertion start position) (Supplementary Table 4). The mate pairs of 6/7 of these Manta SVs mapped to a 541bp region in an intron of *PKD1L1* on chromosome 7. Insertion sequences reported by Severus in chromosome 13 had high sequence similarity to PKD1L1 (Supplementary Table 5). Thus, the insertions in chr13 reported by Severus are concordant with the translocations between chr7 and chr13 reported by Manta. Minda did not deem interchromosomal translocations detected by Manta as concordant with insertions reported by Severus because the second breakend detected by Manta was in a different chromosome to the insertion, i.e. the end positions were not in agreement. This highlights the difficulties regarding SV comparison; determining concordance of further Severus insertions with Manta translocations would require manual inspection of all insertion sequences and breakends. Using ShatterSeek, chromothripsis was not detected between chromosomes 7 and 13 in either short or long read data.

The concordance between SVs detected by Manta and Severus was limited with only 19 (2.4%) and 1 (0.3%) SVs detected in the pre-treatment and post-treatment tumour, respectively (Fig. 4C). A single SV was concordant between Manta and Severus in both the pre- and post-treatment tumours; a 50,484bp deletion in *CDH1* (Fig. 4D). The concordance of SVs detected by Manta was low between the pre- and post-treatment tumours (27/506, 5.3%). Concordant SVs within Manta were predominately breakends (16/19), along with 2 inversions and 1 deletion. Higher concordance was observed within Severus, with 168/447 (37.6%) of SVs called in both the pre- and post-treatment tumours, consisting of 161 insertions and 7 deletions. The high agreement of insertions within Severus is consistent with previous findings (O’Neill et al. 2024).

SV signature analysis was performed. However, insertions are not considered by SigProfiler, along with any deletions, inversions, or duplications smaller than 1kb, resulting in the exclusion of over 80% of SVs detected in tumour samples by Severus. Despite this, some similarity in signature profiles was observed between Manta and Severus with SV2, which is characterised by a high proportion of non-clustered translocations, detected for both callers (Fig. 4E). For Manta, SV4 was detected in the pre-treatment tumour which is attributed to a high proportion of clustered complex translocations. SV signatures detected for Severus were more varied, potentially due to the low amounts of variants considered; SV5 and SV8 were detected in the pre-treatment tumour which are attributed to short (<10kb) deletions and inversions, respectively. SV signatures were developed using short read WGS data. As mentioned, few insertions are detected using short reads and the SVs identified tend to be larger than those detected by long reads. It will be necessary to include SV data from long reads in future SV signature analyses in order to make these signatures more applicable to long reads.

### Methylation patterns

DNA modifications (5-methylcytosine; 5mC and 5-hydroxymethylcytosine; 5hmC) were detected using ONT sequencing. Average rates of DNA modifications were examined across various genomic regions (entire genes, exons, introns, promoters, enhancers, CpG islands, and repetitive regions). Based on genome-wide average CpG modification rates, global hypomethylation was observed in tumours compared to adjacent normal tissues with average global 5mC rates of 70.4% and 72.0% in the pre- and post-treatment tumours and 74.7% and 74.2% in the pre- and post-treatment adjacent normal tissues, respectively. On average, all genomic features were hypomethylated in tumours compared to matched adjacent normal tissues except in CpG islands which were, on average, hypermethylated in tumours (22.0% and 18.5% in pre- and post-treatment tumours, respectively, vs 19.0% and 18.0% in pre- and post-treatment adjacent normal tissues, respectively). Additionally, average global hydroxymethylation rates were similar across samples (2.2%, 2.8%, 3.0% in pre-treatment tumour, pre-treatment adjacent normal and post-treatment adjacent normal, respectively), with the exception of an increased average rate of hydroxymethylation in the post-treatment tumour (4.5%). Higher average rates of hydroxymethylation in the post-treatment tumour were consistent across all genomic regions (Fig. 5A). In post-treatment adjacent normal tissue, the genome-wide methylation rate was lower compared to pre-treatment adjacent normal tissue. This pattern was largely consistent across various genomic features, along with increased average 5hmC rates compared to pre-treatment adjacent normal tissue.

**Figure 5.**
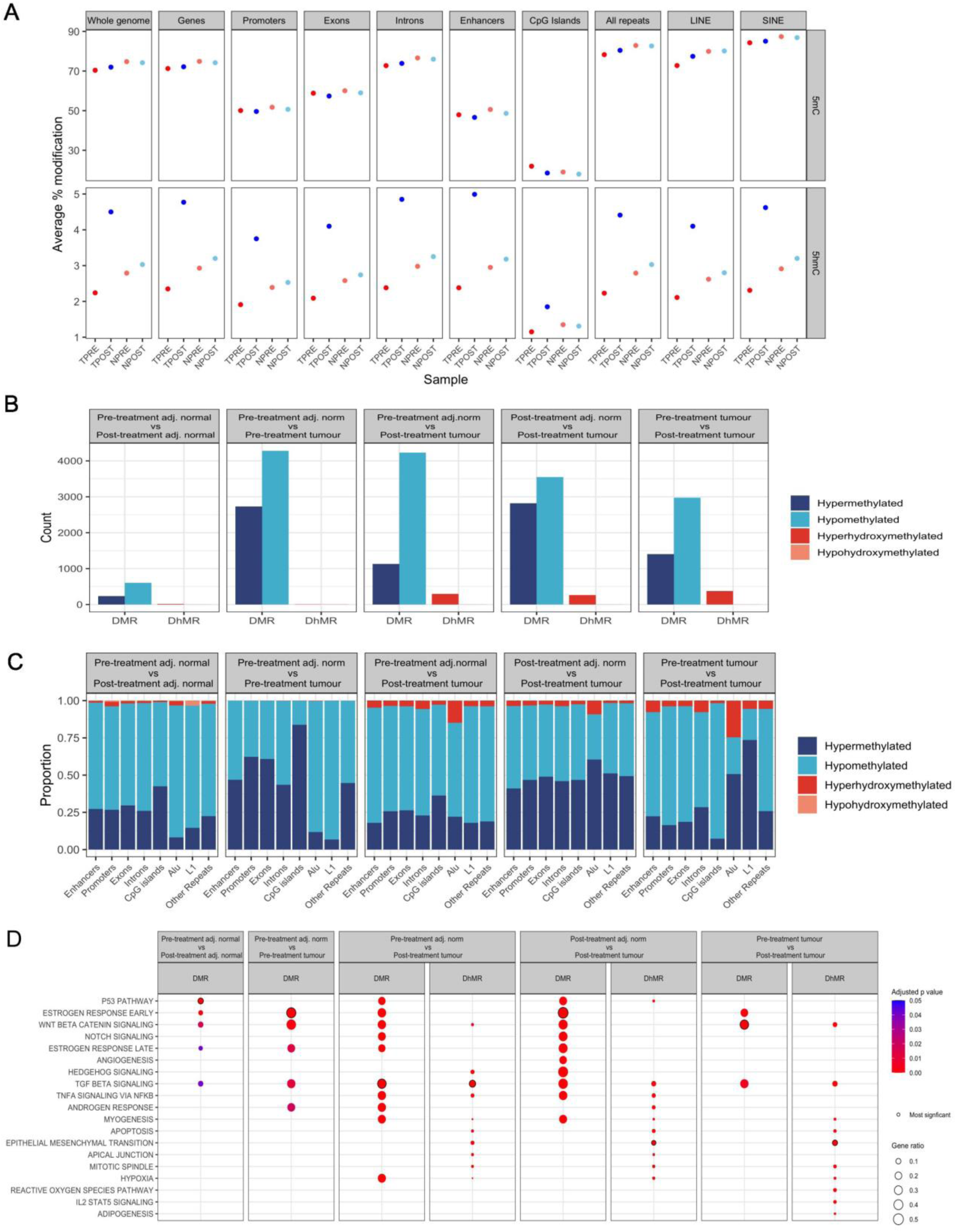
A) Average methylation and hydroxymethylation rates genome-wide and across genomic features. B) Total number of DMRs and DhMRs detected between samples. C) Profiles of differentially modified regions affecting genomic features across sample comparisons. D) Gene set enrichment analysis of D(h)MRs detected across sample comparisons.

Differentially methylated regions (DMRs) and differentially hydroxymethylated regions (DhMRs) were detected between pre- and post-treatment adjacent normal tissues, between tumours and adjacent normal tissues, and between pre- and post-treatment tumours (Fig. 5B). The pre-treatment adjacent normal sample was considered as baseline because the post-treatment adjacent normal had been exposed to chemoradiotherapy. Changes in modification patterns due to chemoradiotherapy can therefore be investigated here. Epigenetic remodelling was more extensive in tumours as fewer DMRs were detected between adjacent normal tissues (839 between pre- and post-treatment adjacent normal tissues) relative to the amount of DMRs detected between adjacent normal tissues and matched tumour samples (7,012 between pre-treatment adjacent normal tissue vs pre-treatment tumour; 5,360 between pre-treatment adjacent normal and post-treatment tumour; 6,372 between post-treatment adjacent normal tissue vs post-treatment tumour) and between pre- and post-treatment tumours (4,378 DMRs). DhMRs were predominately detected in the post-treatment tumour (297 between pre-treatment adjacent normal vs post-treatment tumour; 265 between post-treatment adjacent normal tissue vs post-treatment tumour; 376 between pre-treatment tumour vs post-treatment tumour).

To understand the genomic landscape of DNA modifications, D(h)MRs were annotated with genomic regions. The vast majority of D(h)MRs included multiple genomic features, with a greater number of genomic features of all classes affected by D(h)MRs in tumours. This was in line with the observation of more widespread epigenetic changes (Supplementary Figure 5, Supplementary Table 6). Of all genomic features within D(h)MRs, approximately 25% of enhancers, promoters, exons, and introns were within hypermethylated regions in the post-treatment adjacent normal tissue compared to the pre-treatment adjacent normal tissue (26.7%, 29.6%, 26.0%, 27.3% of promoters, exons, introns, and enhancers respectively) (Figure 5C, Supplementary Figure 5B, Supplementary Table 6). In the pre-treatment tumour compared to the pre-treatment adjacent normal tissue, the proportion of these genomic elements within hypermethylated regions more than doubles for promoters and exons compared to the adjacent normal tissue comparison, with large increases also observed for introns and enhancers (62.3%, 60.8%, 43.5%, 46.8% of promoters, exons, introns, and enhancers respectively). In contrast, the comparison of the pre-treatment adjacent normal to the post-treatment tumour displays a profile more similar to the pre-/post-treatment adjacent normal comparison, with an increased proportion of promoters, exons, introns, and enhancers within hypomethylated regions, but an increased proportion of repetitive elements in hypermethylated regions. In particular, the post-treatment tumour displays an increased proportion of Alu elements in hypermethylated regions but also in hyperhydroxymethylated regions (22.1% in hypermethylated regions and 14.8% in hyperhydroxymethylated regions). A different pattern was observed between the post-treatment adjacent normal sample compared to the post-treatment tumour. An increase in hypermethylated regions was observed across all genomic features. Between pre-treatment tumour and post-treatment tumour, a higher proportion of enhancers, promoters, exons, introns and CpG islands in hypomethylated regions was observed while regions affecting Alu and LINE1 elements were more frequently hypermethylated, with hyperhydroxymethylation occurring across all genomic features.

A pathway enrichment analysis of DMRs detected between the pre-treatment adjacent normal tissue and the post-treatment adjacent normal sample revealed several cancer-related pathways (Fig 5D). The most significantly enriched pathway in was the P53 pathway, potentially reflecting epigenetic changes affecting the DNA damage response and cellular survival mechanisms due to damage inflicted by neoadjuvant chemoradiotherapy. The P53 pathway was also enriched in the post-treatment tumour compared to adjacent normal tissues, but not in the pre-treatment tumour. Compared to timepoint-matched adjacent normal tissues, DMRs in the pre-treatment and post-treatment tumours were most significantly enriched for genes involved in the early estrogen response which has been posited to play a role in CRC progression (Caiazza et al. 2015). TGFβ signalling was significantly enriched in all comparisons, though it was the most significant pathway identified for DMRs and DhMRs detected between the pre-treatment adjacent normal tissue and post-treatment tumours. TGFβ signalling has been shown to promote epigenetic changes that induce epithelial-mesenchymal transition (EMT) (Kim et al. 2020). Moreover, DhMRs detected between the pre- and post-treatment tumours were most significantly enriched for the EMT pathway. EMT promotes cancer cell stemness through epigenetic reprogramming, increasing metastatic potential and inducing chemotherapy resistance (Kim et al. 2020). The Wnt β-catenin pathway was most significantly enriched in DMRs detected between pre-treatment and post-treatment tumours. This pathway is also a key promoter of EMT and has been shown to induce 5-fluorouracil resistance in CRC (Xue et al. 2024; Dong et al. 2022). This patient developed metastatic disease affecting the liver and omentum 9 months after beginning nCRT.

Overall, differential methylation analysis showed changes in methylation patterns that are commonly observed in tumours, including global hypomethylation and local hypermethylation of enhancers, promoters and CpG islands (Nishiyama and Nakanishi 2021). This, along with hypomethylation of Alu and LINE1 elements in tumours compared to adjacent normal tissues illustrates an altered epigenetic landscape in tumours. D(h)MRs in tumours affected genes involved in signalling through the TGFβ and Wnt β-catenin pathways, both of which are associated with EMT. However, epigenetic remodelling also occurred in normal tissues to a lesser extent; in the post-treatment normal tissue, DMRs were enriched for genes involved in p53 signalling suggesting that this pathway was induced in response to DNA damage inflicted by chemoradiation (Kong et al. 2021).

### Potential driver mutations

Potential driver mutations in *KRAS* (p.G12D), *PIK3CA* (p.E542K) and *SOCS1* (p.Q131*) were consistently detected by short and long reads in the pre-treatment tumour (Table 3). The *KRAS* variant was also successfully called in the short and long read post-treatment tumour. However, detection of *PIK3CA* and *SOCS1* drivers in the post-treatment tumour was inconsistent. Though present at a VAF of 20%, the *PIK3CA* variant was not successfully called in the long read post-treatment tumour as it was also present at 2.9% VAF in the long-read germline sample. The variant was only supported by a single read in ONT WGS of the germline sample and was not detected in the short-read germline sample. Moreover, this germline VAF was only reported in the post-treatment tumour iteration of ClairS and not the pre-treatment tumour iteration – both of which used the same germline input. This highlights a discrepancy in the ClairS somatic calls at this position between pre- and post-treatment tumour iterations. The *SOCS1* variant was not initially reported in the post-treatment tumour sequenced by short reads. However, force-calling with Mutect at that position detected evidence of the variant (VAF 4.2%) but FilterMutectCalls deemed this variant a likely sequencing error.

**Table 3.**
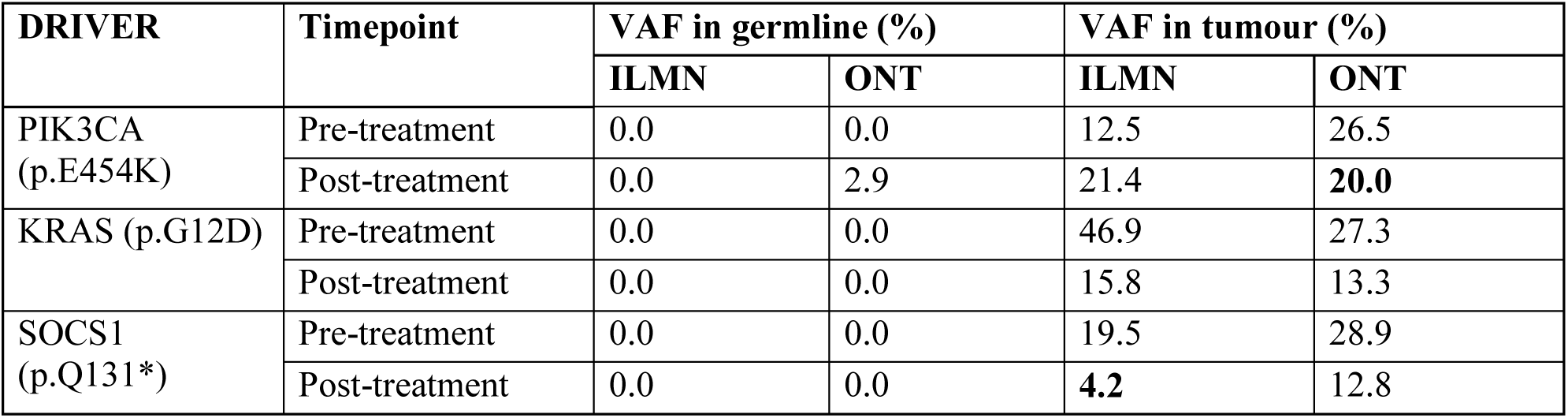
Driver SNVs detected by short and long read sequencing. Bold lettering indicates that the mutation did not pass quality thresholds in the technology indicated.

Potential driver CNAs were largely consistent; with the exception of several false positive homozygous deletions reported in the long reads due to limitations with the copy number caller utilised. Loss of heterozygosity in potential drivers including CDK12, CTNNB1, ERRB2, TP53 and USP36 was consistently detected by short and long reads.

Few structural variants were consistently detected between short and long read technologies; only a deletion in *CDH1* was detected by long and short reads in both pre- and post-treatment tumours. The only insertion detected in *CTNNB1* by short reads in the pre-treatment tumour was also confirmed by long reads in the pre- and post-treatment tumour (Table 4). Interestingly, both of these genes have roles in EMT (Busch et al. 2017). Moreover, *CTNNB1* encodes β-catenin which is a crucial component of the Wnt signalling pathway, mutations in which are known to drive CRC tumourigenesis (Liu et al. 2022). A 136bp insertion in *CTNNB1* was detected by short and long reads in the pre-treatment tumour, and by long reads in the post-treatment tumour but not short reads. Additionally, LOH of *CTNNB1* was detected by long and short reads.

**Table 4:**
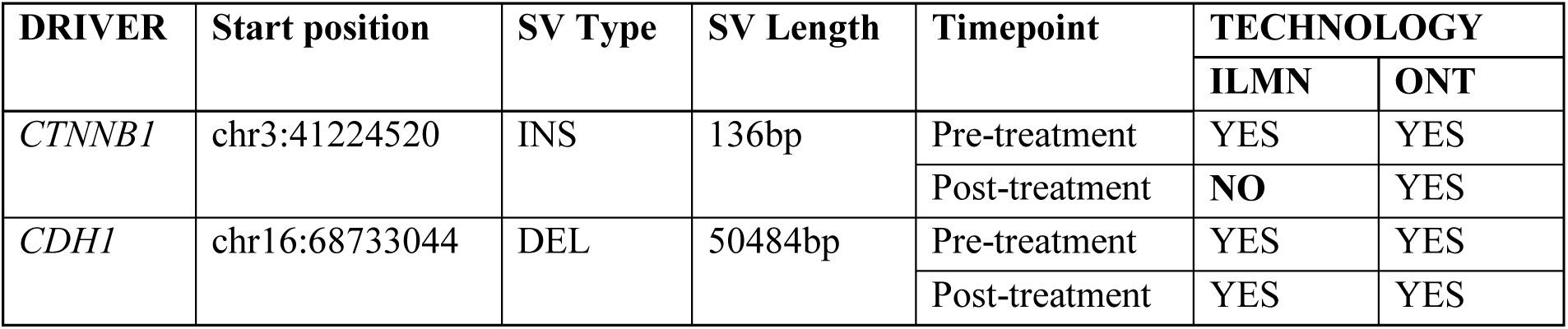
Driver SVs detected by short and long read sequencing. Bold lettering indicates that the mutation was not called in the technology indicated.

*CDH1* is a tumour suppressor gene that encodes E-cadherin, a regulator of β-catenin and suppressor of EMT. A 50,484bp deletion in *CDH1* was detected in both pre- and post-treatment tumours by short and long reads. The 50kb deletion encompasses approximately 4.2kb upstream of exon 1, the entirety of exons 1 and 2, and extends into intron 2, disrupting the structure of this tumour suppressor gene. Exons 1 and 2 encode the signal peptide of E-cadherin which is necessary for the translation and translocation of this protein, with mutations affecting the signal peptide linked to hereditary diffuse gastric cancer (Figueiredo et al. 2018; Selvanathan et al. 2020; Wang et al. 2004).

Epigenetic modification of 61 driver genes was assessed. The list of drivers was chosen based on the COSMIC cancer gene census and genes with potential driver mutations identified in the tumours in this study were also added. Of these genes, 46 were affected by D(h)MRs in at least one comparison. Enhancers linked to driver genes were the predominant genomic feature encompassed by D(h)MRs, with fewer promoters and gene bodies affected (Figure 6). Within gene bodies, introns were more often affected by D(h)MRs than exons. In terms of specific drivers, the promoter, exons and introns of FBLN2, and RSPO3 were affected by D(h)MRs, as well as enhancer regions associated with these genes, in the pre-treatment tumour compared to the pre-treatment adjacent normal sample. The same pattern was not observed between the post-treatment tumour and the pre-treatment adjacent normal sample.

**Figure 6.**
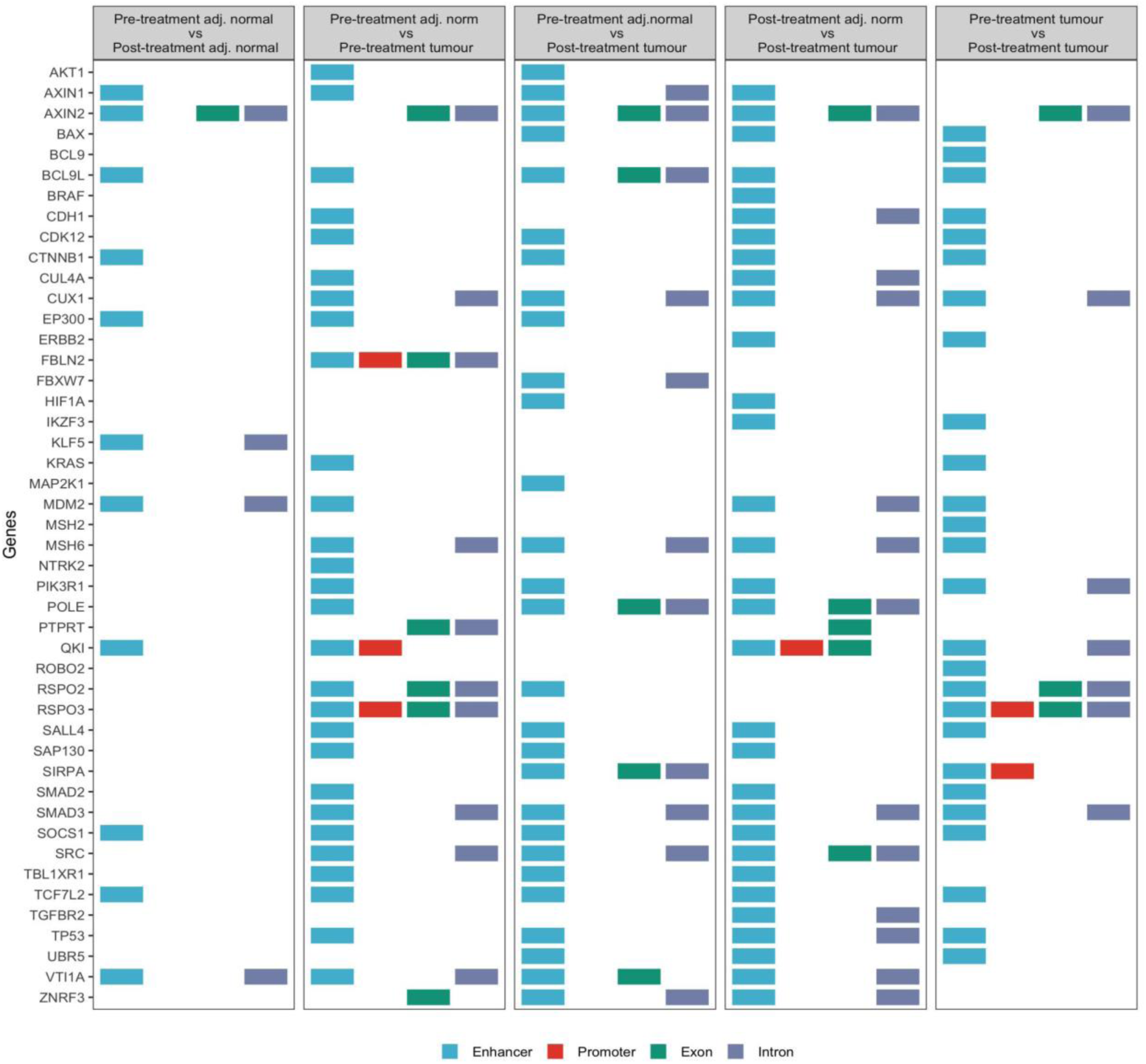
Regions of driver genes affected by differentially modified regions.

## DISCUSSION

Here we compared somatic variants detected in ILMN and ONT WGS data derived from pre- and post-treatment tumour samples of a locally advanced rectal cancer patient. SNV concordance between technologies was significantly affected by tumour and germline sample purity, while indel concordance was expectedly poorer. Differences in the reporting of structural variants made comparison challenging even when using an SV agnostic comparison tool. Copy number profiles were largely consistent between ILMN and ONT. Methylation data available from ONT WGS data provided added further context to the tumour profiles and the effect of nCRT on the epigenomic landscape.

We found that the majority of SNVs could be detected by both short and long read technologies, but short reads were not as capable of detecting SNVs in repetitive regions. High purity samples are important when detecting somatic mutations as variant calling algorithms consistently show poorer performance when detecting low frequency variants, especially at lower tumour purities (Zheng et al. 2023; Chen et al. 2020). Here, the importance of high purity samples was evident because low tumour purity of the post-treatment tumour significantly affected SNV concordance in both short and long read data. Notably, purity of normal samples also affects the performance of variant calling tools (Zheng et al. 2023). Discrepancies in detecting driver SNVs were observed in our study. The *PIK3CA* E545K mutation was not initially detected in the long read post-treatment tumour despite its presence at a VAF of 20.0%.

Accurate and reliable indel detection is a challenging task. Somatic variant callers consistently display markedly poorer performance for indel detection compared to SNV detection and the concordance of indel calls is limited between variant callers (Zheng et al. 2023; Park et al. 2024; Bennett et al. 2020; O’Rawe et al. 2013). It is estimated that approximately one third of indels in the human genome remain undetected, for example due to the difficulty calling indels in repetitive sequence such as homopolymers (Bennett et al. 2020; Narzisi and Schatz 2015). Here we report limited indel concordance between short and long read technologies. Across all samples, an average of 66.1% of indels detected were within or adjacent to simple homopolymers, with most discordant indels occurring within/adjacent to simple homopolymers. This is consistent with previous work highlighting homopolymers as difficult regions for indel calling (Gustafson et al. 2024).

Generally, copy number calls across the genome were consistent between short and long reads. However, copy number calls for specific driver genes were less congruous; homozygous deletions of several driver genes were reported in long read but not short read WGS data.

A main advantage of long reads is the ability to detect smaller structural variants as well as complex structural variants. Similar to previous work, we observed that Severus detected a greater number of insertions and smaller deletions compared to Manta (Keskus et al. 2025). We found that concordance of structural variants was low between technologies; however, this was exacerbated by differences in how structural variant calling tools report variants. While short read tools report individual breakpoints, long reads can encompass entire translocations and report them as insertions. Therefore, assessment of structural variant concordance by comparing genomic coordinates was not straightforward; several insertions reported by Severus were reported as translocations by Manta which resulted in false negative concordance calls.

Mutational signature analysis provides insight into the biological processes acting in a tumour. SNVs, indels, CNAs, and SVs have been assigned to specific mutational profiles and associated with exogenous and/or endogenous mutagens (Alexandrov et al. 2020; Steele et al. 2022; Li et al. 2020). Mutations included in these profiles have thus far been derived from short read sequencing data. Here we sought to assess the relevance of these established short-read-based mutational signatures to long read mutational profiles. Single base substitution signatures were consistently detected by short and long reads in pre- and post-treatment tumours which indicates that similar biological processes reflected by SNV patterns can be detected using short and long read sequencing data. Inconsistencies in indel signatures were indicative of low indel concordance rather than unsuitability of indel signatures for long read data. Copy number signatures were largely consistent between short and long read data though a signature associated with sequencing artefacts was detected in the long read data. Finally, structural variant signatures were not suitable for use with long read data as insertions and other SVs shorter than 1kb are not accepted as input for structural variant signature analysis using SigProfiler. We note that a limitation of this study is the assessment of one variant calling tool per mutation type.

Tumour evolution can be monitored by tracking somatic mutations at various timepoints. In this study, pre- and post-treatment tumour samples were available. Subclones carrying mutations that confer a selective advantage will outcompete other subclones. As mentioned, tumour purity is crucial to accurate variant calling. In terms of capturing tumour evolution, this poses a challenge when post-treatment tumours may be less pure if the patient has partially responded to treatment or due to sampling issues. Reduced concordance of SNVs and indels between pre-treatment tumour and post-treatment tumour may reflect the lower purity of the post-treatment tumour as well as the loss of subclones contributing to tumour heterogeneity. In terms of CNAs, copy number profiles were generally stable except for a loss detected in chromosome 12 by both short and long reads. For short reads, most structural variants were detected in the pre- or post-treatment tumour only. While this was also true for long reads, many SVs were still concordant between the pre- and post-treatment tumour.

The genomic characteristics of the locally advanced rectal cancer genome studied here were consistent between short and long reads. The tumour was histologically confirmed as MSS; consistent with this, no observations of MSI were made in short or long read data. *KRAS* G12D and *PIK3CA* E545K mutations were detected in the tumour genome by short and long read WGS data. *KRAS* G12D and *PIK3CA* E454K mutations occur in 11.8% and 5.4% of MSS stable CRCs, respectively (Cornish et al. 2024). Approximately 50% of non-hypermutated early onset CRCs display early *TP53* mutations with whole genome doubling (Kim et al. 2021). Consistent with this, in this 37-year-old patient we observed a non-hypermutated tumour genome and LOH of *TP53* with a subsequent WGD event in both short and long read data. SVs affecting *CTNNB1* and *CDH1*, both of which are involved in the Wnt pathway, were consistently supported by short and long read data. A 136bp insertion was detected in intron 2 of *CTNNB1* and was predicted to result in the loss of the acceptor site (probability 0.89). GSK-3β is a regulator of β-catenin and phosphorylates sites within exon 3 of β-catenin to promote its degradation (Gao et al. 2017). Hotspot mutations within exon 3 of β-catenin stabilise the protein, leading to greater signalling through the Wnt signalling pathway to drive tumourigenesis (Kim and Jeong 2019). In-frame deletions resulting in the complete loss of exon 3 have been shown to enable activation of the Wnt signalling pathway, with this type of *CTNBB1* mutation occurring in multiple cancer types, but most frequently in CRC (Yaeger et al. 2018). As the 136bp insertion detected here affects transcript splicing, it is possible that exon 3 is skipped in this tumour to produce a more stable isoform of β-catenin. The Wnt β-catenin pathway is a key driver of EMT and can induce 5-FU resistance in CRC (Xue et al. 2024; Dong et al. 2022). The additional methylation data from ONT WGS provided further context as to how Wnt pathway signalling and other EMT-related pathways were affected by nCRT in this tumour which provides potential further insight into the patient’s poor response to nCRT and subsequent development of metastasis.

Overall, key characteristics of this genome were captured by both sequencing technologies, with ONT DNA modification data providing additional insights. Mutational signatures for small somatic variants appear to be useful for interpretation of somatic SNVs but are less applicable to CNVs and SVs detected with long reads. Computational tools for somatic variant calling with long read WGS data are in early stages of development relative to short-read tools and thus care should be taken when applying these tools. However, long read ONT WGS shows great utility in characterising cancer genomes.

## METHODS

### Clinical samples

Informed consent was provided by the patient to perform whole genome sequencing (WGS) and ethical approval was obtained for the study (reference: 13/82 and 15/09). In total, 5 samples were collected from the patient. At baseline (pre-treatment), a pre-treatment primary tumour sample and adjacent normal tissue were obtained, as well as matched blood. During surgical resection, a post-treatment primary tumour sample and adjacent normal tissues were collected.

### DNA Extraction

High molecular weight DNA was extracted from all samples following the Oxford Nanopore Rabbit Liver DNA extraction protocol, with one modification due to small sample size; the Blood and Cell Culture DNA Mini Kit (QIAGEN) was used instead of the Blood and Cell Culture DNA Midi Kit (QIAGEN). DNA was quantified using the Qubit dsDNA Broad Range Assay Kit (Invitrogen). DNA quality was determined using the Bioanalyser DNA 1000 kit (Agilent Technologies).

### Oxford Nanopore whole genome sequencing

DNA libraries were prepared using the Ligation Sequencing Kit (SQK-LSK114, ONT), as per the manufacturer’s protocol. Sequencing was performed with R10.4.1 PromethION flow cells (FLO-PRO144M) by a PromethION 2 Solo machine with a runtime of 72 hours per flowell. Each flowcell was washed and reloaded as was necessary based on pore performance. Live basecalling was performed using the MinKNOW software (v23.11.4) with Dorado (v7.2.13) on high accuracy mode with 5mC and 5hmC calling enabled. Reads with a quality score less than 7 were discarded. FASTQ files were aligned to GRCh38 using minimap2 (v2.26) with the following settings; -ax map-ont (Li 2021). BAM files were sorted and indexed using samtools (v1.9) (Danecek et al. 2021). Genome coverage was determined using mosdepth (v0.3.3) (Pedersen and Quinlan 2017). Somalier (v0.2.19) was used to ensure samples were related (Pedersen et al. 2020).

### Illumina whole genome sequencing

Extracted DNA was diluted using elution buffer. At Novogene, DNA libraries were prepared and paired-end sequencing carried out on an Illumina NovoSeq 6000. Adapter trimming was performed on FASTQ files using Trimmomatic (v0.39) with a modified TruSeq2-PE.fa file to remove partial adapter contamination (Supplementary Materials File 1) (Bolger et al. 2014). The parameters were ILLUMINACLIP:<adapters.fa>:4:4:5:2:True LEADING:3 TRAILING:3 MINLEN:36. Trimmed FASTQ files were aligned to GRCh38 using BWA-mem (v0.7.17) (Li 2013). Using samtools (v1.12) (Danecek et al. 2021), BAMs were sorted and where samples were sequenced across multiple lanes, BAM files were merged prior to marking duplicates. Samtools was used to index BAM files. Using GATK (v4.5.0.0), duplicate reads were marked (MarkDuplicates) and base quality score recalibrated (BaseRecalibrator, ApplyBQSR) (McKenna et al. 2010). Genome coverage was assessed using mosdepth (v0.3.3) (Pedersen and Quinlan 2017). Somalier (v0.2.19) was used to confirm sample relatedness (Pedersen et al. 2020).

### Small somatic variant calling

Small somatic variants were detected in Illumina WGS by Mutect2 (GATK v.4.3.0.0) using by comparing tumour/adjacent normal tissues to matched blood. A panel of normals from the 1000 Genomes Project (1000g_pon.hg38.vcf.gz) and a germline resource from gnomAD (af-only-gnomad.hg38.vcf.gz) were supplied to Mutect2 to reduce false positive somatic calls. The ExAC common germline variants (small_exac_common_3.hg38.vcf.gz) were used with GATK (v4.3.0.0) GetPileupSummaries and CalculateContamination. Outputs from these tools were used in FilterMutectCalls to filter technical artefacts, non-somatic mutations, and sequencing errors. After these filtering steps, the VCF was split into SNVs and indels using bcftools view.

Using ClairS (v0.4.0), single nucleotide variants and indels were detected in ONT WGS data by comparing tumour/adjacent normal tissues to matched blood (Zheng et al. 2023). The following parameters were set; --platform ont_r10_dorado_hac_5khz --enable-indel-calling --enable_clair3_germline_output –q8 –longphase_for_phasing. Enabling germline output was necessary to obtain germline variant VCFs required as input for other tools (i.e. Wakhan and Severus).

Somatic small variants were required to meet the following criteria; within chromosomes 1-22, X and Y, marked as PASS, and with read depths of at least 4 in both the tissue of interest and the germline sample. Multiallelic variants were split using bcftools norm and indels normalised using GATK (v4.3.0.0) LeftAlignAndTrimVariants. Indels over 50bp long were removed. Unique variant IDs in the format of ‘<chromosome>:<position>:<reference_allele>:<alternate_allele>’ were assigned to each variant using bcftools annotate. Variant effect predictor (VEP; v.107) was used to annotate small somatic variants (McLaren et al. 2016). To remove common variants, filter_vep was used to remove variants that were not listed in the Catalogue of Somatic Mutations in Cancer (COSMIC) database (v95) or were not novel variants (--filter “(Existing_variation match COS or not Existing_variant)”). Variant allele frequency was calculated as read depth of alternate allele/total read depth reported by a given caller.

### Small variant concordance

SNVs were considered concordant between sequencing technologies on the basis of SNV IDs; the chromosome, position, reference and alternate allele must agree. To evaluate indel concordance, several bedtools (v2.30.0) approaches were assessed; 1) complete and reciprocal overlap of indels (--f 1, -r), 2) minimum overlap of 1bp (default settings), and 3) 1bp overlap within a 5bp window (slop –b 5) (Quinlan and Hall 2010). Similar bedtools approaches were used to assess overlap of indels with homopolymers. Simple repeat homopolymer files (GRCh38_SimpleRepeat_homopolymer_*_slop5.bed.gz) were obtained from Genome in a Bottle genome stratifications (https://github.com/usnistgov/giab-stratifications) (Dwarshuis et al. 2024).

Factors affecting SNV concordance were examined in further detail. Variants detected only in short-read WGS data by Mutect2 were evaluated in ONT data through another iteration of ClairS in which an additional options were used as follows: --genotyping_mode_vcf_fn was used to supply a VCF with SNVs private to Mutect and –phase_tumor to re-activate phasing of ONT data when supplying a VCF. Variants detected only in ONT WGS data were evaluated in ILMN data by using Mutect2 to force call at the SNV positions by supplying a VCF with SNVs private to ClairS to the –alleles option. Using the annotatr R package (v1.24.0), SNVs were annotated with genomic region (“hg38_genes_1to5kb”, “hg38_genes_exons”, “hg38_genes_promoters”, “hg38_genes_introns”, “hg38_genes_intergenic”). SNVs were annotated with repetitive regions using RepeatMasker information for hg38 available from the AnnotationHub R package (v3.6.0).

### Structural Variant calling

Structural variants in Illumina WGS data were detected using Manta (v1.6.0) with default parameters (Chen et al. 2016). The supplementary convertInversion.py script was applied to the output VCF to reformat inversion entries to single inverted sequence junctions.

Structural variants in ONT WGS data were detected using Severus (v1.3) (Keskus et al. 2025). BAMs were phased using WhatsHap (v1.7) haplotag (Martin et al. 2016). Phased germline SNVs output by ClairS were provided to the haplotag command to ensure consistent phasing between samples. Phased tumour and germline BAMs and the phased germline VCF were used as input for Severus. Tandem repeat regions (human_GRCh38_no_alt_analysis_set.trf.bed.gz) were supplied to Severus using –vntr-bed.

For Manta and Severus, structural variants marked as PASS and within chromosomes 1:22, X, and Y were selected for further analysis. Comparison of structural variants as performed using Minda (https://github.com/KolmogorovLab/minda, commit 2066ef5) with default parameters. Circos plots were generating using the circlize R package (v0.4.15). The effect of structural variants on splicing was investigated using the online version of SpliceAI (https://spliceailookup.broadinstitute.org/, accessed 31/07/2025) (Jaganathan et al. 2019). ShatterSeek (v1.1) was used to detect chromothripsis.

### Somatic Copy Number Alterations

facetsSuite (https://github.com/mskcc/facets-suite, v2.0.9) was used to run FACETS (v0.6.2) on the Illumina WGS data to detect sCNAs. FACETS requires a base-level coverage file as input which must be generated using snp-pileup (v0.6.2). The parameters -q15 -Q20 -P100 -r25,0 were applied as well as a file outlining common SNPs (obtained from dbSNP, build 151). The resulting pileup was used as input to the facetsSuite script run_facets_wrapper.R.

Wakhan (https://github.com/KolmogorovLab/Wakhan, v0.1.0) was used to detected somatic copy number alterations in ONT WGS data. BAMs haplotagged with using phased normal VCFs output from ClairS were used as input. The normal phased VCF from ClairS and somatic SV breakpoints from Severus were provided to Wakhan using the –normal-phased-vcf and –breakpoints options, respectively. Subclonal analysis (--copy-numbers-subclonal-enable) and detection of loss of heterozygosity (--loh-enable) were enabled. A purity range was provided (0-0.60). For comparative purposes, a list of all genes assessed by FACETS was provided to Wakhan using user-input-genes. Copy number profiles were visualised using R.

### Mutational Signature Analysis

Single base substitutions, indels, copy number, and structural variant signatures were assigned using SigProfilerAssignment (v0.1.8) with COSMIC v3.4 (Islam et al. 2022). To perform copy number analysis on ONT data, haplotype-specific bed files output by Wakhan were combined using awk and modified to replicate the output format of the FACETS segmentation file. The version of Wakhan used in this analysis (v0.1.0) does not perform copy number calling in centromeric regions and as a result segments surrounding centromeres reported 0 coverage. Segments flanking centromeres were extended to fill in centromeric gaps and avoid introduction of false homozygous deletions to the copy number signature analysis. For structural variant signature analysis, output VCFs of Manta and Severus were filtered for PASS variants and converted to BEDPE format. All mutational signature contributions and mutational matrices were visualised in R.

### Methylation analysis

Unmapped BAMs containing methylation information were output by the MinKNOW software. Unmapped BAMs were sorted by read name and merged. Merged uBAMs were converted to FASTQ format using –T * to ensure all tags were carried over and subsequently aligned to GRCh38 using minimap2 (v2.26) with the –ax map-ont –y options. Aligned BAMs were sorted by coordinate and reads with MAPQ < 20 were removed. Longphase (v1.7.3) haplotag was used to phase methylBAMs using germline SNVs detected by ClairS and SVs detected by Sniffles (v2.6.0) (Lin et al. 2022; Smolka et al. 2024). Modkit (https://github.com/nanoporetech/modkit, v0.5.0) pileup was used to generate bedmethyl files. Bedmethyl files were filtered for all sites within autosomes and sex chromosomes. The R package DSS (v 2.54.0) was used to detect differentially modified regions between samples. DMLtest() was used with smoothing enabled as a single replicate analysis was performed. CallDMR() was used with minlen = 100, minCG = 15, dis.merge = 100. The minimum change in modification (delta) was set to 15% for DMRs and 5% for DhMRs. Differentially modified regions were linked to genomic features using bedtools intersect. The genomic coordinates of canonical transcripts (knownGene, exons, introns) for GRCh38 were obtained from UCSC table browser. Promoters were defined as –2Kb and +200bp from the transcription start site. RepeatMasker and CpG island coordinates for GRCh38 were also obtained from UCSC table browser. Enhancers were obtained from GeneHancer v5.24 (Fishilevich et al. 2017). Only elements with elite status classed as enhancers or enhancer/promoters were included in this analysis. Functional enrichment of genomic regions was performed using rGREAT (v2.9.2) with windows extending from genes and the UCSC knownGene transcription start sites (Gu and Hübschmann 2023). Hallmark MSigDB gene sets were used.

## DATA ACCESS

The genomic datasets generated in this study will be available from European Genome-Phenome Archive for academic use through the Genomic Oncology Research Group Data Access Committee (EGAC00001001585).

## AUTHOR CONTRIBUTIONS

L.M., data analysis; L.M., bioinformatic data processing; B.ON., J.F., L.OS., A.M., L.G., D.MM., B.T.H., S.T., clinical data and sample acquisition;; L.M, S.T., ONT sequencing, L.M, wrote the manuscript; critical revision an approval of final manuscript (all authors); B.T.H, S.J.F, S.T., study supervision. S.J.F study design and concept.

## ACKNOWLEDGEMENTS

We would like to thank the patient for kindly providing tumour tissue to enable this research study. We would like to thank the Irish Centre for High End Computing (https://www.ichec.ie/) for the use of HPC infrastructure.

