## Supplemental material for "Whole genome sequencing of pre-treatment and post-treatment locally advanced rectal cancer using long and short read technologies"

**Contents:**

**Supplementary Materials: Modified TruSeq2 paired end adapter FASTA file**

**Supplementary Tables 1-6**

**Supplementary Figures 1-5**

### Supplementary Materials: Modified TruSeq2 paired end adapter FASTA file

```
> PrefixPE/1
AATGATACGGCGACCACCGAGATCTACACTCTTTCCCTACACGACGCTCTTCCGATCT
> PrefixPE/2
CAAGCAGAAGACGGCATACGAGATCGGTCTCGGCATTCCTGCTGAACCGCTCTTCCGATCT
> PCR_Primer1
AATGATACGGCGACCACCGAGATCTACACTCTTTCCCTACACGACGCTCTTCCGATCT
> PCR_Primer1_rc
AGATCGGAAGAGCGTCGTGTAGGGAAAGAGTGTAGATCTCGGTGGTCGCCGTATCATT
> PCR_Primer2
CAAGCAGAAGACGGCATACGAGATCGGTCTCGGCATTCCTGCTGAACCGCTCTTCCGATCT
> PCR_Primer2_rc
AGATCGGAAGAGCGGTTCAGCAGGAATGCCGAGACCGATCTCGTATGCCGTCTTCTGCTTG
> FlowCell1
TTTTTTTTTTAATGATACGGCGACCACCGAGATCTACAC
> FlowCell2 TTTTTTTTTTCAAGCAGAAGACGGCATACGA
> PartialInsertion1
CAGAAGACGGCATACG
> PartialInsertion2
AGAAGACGGCATACG
> PartialInsertion3
GAAGACGGCATACG
> PartialInsertion4
AAGACGGCATACG
> PartialInsertion5
AGACGGCATACG
> PartialInsertion6
GACGGCATACG
> PartialInsertion7
ACGGCATACG
> PartialInsertion8
CGGCATACG
> PartialInsertion9
GGCATACG
> PartialInsertion10
GCATACG
> PartialInsertion11
CATACG
> PartialInsertion1_rc
CGTATGCCGTCTTCTG
> PartialInsertion2_rc
CGTATGCCGTCTTCT
> PartialInsertion3_rc
CGTATGCCGTCTTC
> PartialInsertion4_rc
CGTATGCCGTCTT
> PartialInsertion5_rc
CGTATGCCGTCT
> PartialInsertion6_rc
CGTATGCCGTC
> PartialInsertion7_rc
CGTATGCCGT
> PartialInsertion8_rc
CGTATGCCG
> PartialInsertion9_rc
CGTATGCC
> PartialInsertion10_rc
CGTATGC
> PartialInsertion11_rc
CGTATG
```

**Supplementary Table 1:** Average ILMN and ONT WGS coverage per sample.

| Sample | ILMN | ONT |
|---:|:---:|:---:|
| Blood | 42X | 32X |
| Pre-treatment tumour | 56X | 48X |
| Post-treatment tumour | 41X | 45X |
| Pre-treatment adjacent normal | 42X | 41X |
| Post-treatment adjacent normal | 44X | 20X |

**Supplementary Table 2:** Total SNVs detected in tumours and SNVs with all available information for regression.

| | Total count | Mutect/ClairS intersection | Private to Mutect | Private to ClairS |
|---|---|---|---|---|
| Total SNVs detected in tumours | 16,216 | 8,432 | 4,747 | 3,037 |
| SNVs validated | 15,605 | 8,432 | 4,574 | 2,599 |
| Orthogonal data/caller could not validate SNV | 611 | 0 | 173 | 438 |

**Supplementary Table 3:** Counts of indels overlapping or within 5bp of simple homopolymers by caller and sample.

| Caller | Variant | Category | Tumour (% calls) | | Adjacent normal (% calls) | |
|---|---|---|---|---|---|---|
| | | | Pre-treatment | Post-treatment | Pre-treatment | Post-treatment |
| **Mutect** | Indels | Total | 672 (100) | 350 (100) | 124 (100) | 117 (100) |
| | | Overlapping/within HP | 428 (63.7) | 195 (55.7) | 78 (62.9) | 77 (65.8) |
| | | Not overlapping/within HP | 244 (36.3) | 155 (44.3) | 46 (37.1) | 40 (34.2) |
| | Insertions | Total | 237 (100) | 96 (100) | 19 (100) | 45 (100) |
| | | Overlapping/within HP | 184 (76.7) | 64 (66.7) | 8 (42.1) | 27 (60.0) |
| | | Not overlapping/within HP | 53 (23.3) | 32 (33.3) | 11 (57.9) | 18 (40.0) |
| | Deletions | Total | 435 (100) | 254 (100) | 105 (100) | 72 (100) |
| | | Overlapping/within HP | 244 (56.1) | 131 (51.6) | 70 (66.7) | 50 (69.4) |
| | | Not overlapping/within HP | 191 (43.9) | 123 (48.4) | 35 (33.3) | 22 (30.6) |
| **ClairS** | Indels | Total | 714 (100) | 532 (100) | 378 (100) | 606 (100) |
| | | Overlapping/within HP | 451 (63.2) | 368 (69.2) | 277 (73.3) | 449 (74.1) |
| | | Not overlapping/within HP | 263 (36.8) | 164 (30.8) | 101 (26.7) | 157 (25.9) |
| | Insertions | Total | 270 (100) | 228 (100) | 162 (100) | 295 (100) |
| | | Overlapping/within HP | 166 (61.5) | 139 (61.0) | 96 (59.3) | 179 (60.7) |
| | | Not overlapping/within HP | 104 (38.5) | 89 (39.0) | 66 (40.7) | 116 (39.3) |
| | Deletions | Total | 444 (100) | 304 (100) | 216 (100) | 311 (100) |
| | | Overlapping/within HP | 285 (64.2) | 229 (75.3) | 181 (83.8) | 270 (86.8) |
| | | Not overlapping/within HP | 159 (35.8) | 75 (24.7) | 35 (16.2) | 41 (13.2) |

**Supplementary Table 4:** Insertions detected by Severus in chromosome 13 showed concordance with Manta BNDs with mates in chromosomes 4 and 7.

| Sample | Severus | | Manta | | | |
|---|---|---|---|---|---|---|
| | **ID** | **CHROM:POS** | **BND_1** | **CHROM:POS** | **BND_2** | **CHROM: POS** |
| Pre-treatment tumour | INS10452 | chr13:46267547 | MantaBND:6:7196:23139:0:1:0:1 | chr13:46267543 | MantaBND:6:7196:23139:0:1:0:0 | chr7:47825323 |
| | INS10474 | chr13:53792439 | MantaBND:6:7196:23150:0:1:0:0 | chr13:53792440 | MantaBND:6:7196:23150:0:1:0:1 | chr7:47825430 |
| | | | MantaBND:6:7196:23150:1:0:0:1 | chr13:53792533 | MantaBND:6:7196:23150:1:0:0:0 | chr7:47825830 |
| | INS10486 | chr13:57288897 | MantaBND:6:40870:40871:0:0:0:1 | chr13:57288898 | MantaBND:6:40870:40871:0:0:0:0 | chr4:15869900 |
| | INS10502 | chr13:61258367 | MantaBND:6:7196:23129:0:0:0:0 | chr13:61258382 | MantaBND:6:7196:23129:0:0:0:1 | chr7:47825664 |
| | INS10518 | chr13:65342825 | MantaBND:6:7196:23138:0:0:0:1 | chr13:65342768 | MantaBND:6:7196:23138:0:0:0:0 | chr7:47825620 |
| | | | MantaBND:6:7196:23138:1:0:0:0 | chr13:65342868 | MantaBND:6:7196:23138:1:0:0:1 | chr7:47825204 |
| | NS10559 | chr13 78481529 | MantaBND:6:7196:23166:1:0:0:0 | chr13:78481392 | MantaBND:6:7196:23166:1:0:0:1 | chr7:47825953 |
| | | | MantaBND:6:7196:23166:0:1:0:0 | chr13:78481545 | MantaBND:6:7196:23166:0:1:0:1 | chr7:47825566 |
| | INS10635 | chr13:92931746 | MantaBND:6:7196:23161:0:5:0:0 | chr13:92931750 | MantaBND:6:7196:23161:0:5:0:1 | chr7:47825559 |
| | | | MantaBND:6:7196:23161:1:0:0:0 | chr13:92931818 | MantaBND:6:7196:23161:1:0:0:1 | chr7:47825967 |
| Post-treatment tumour | INS10584 | chr13:46267547 | MantaBND:5:18348:18365:1:0:0:1 | chr13:46267394 | MantaBND:5:18348:18365:1:0:0:0 | chr7:47825889 |
| | | | MantaBND:5:18348:18365:0:0:0:1 | chr13:46267543 | MantaBND:5:18348:18365:0:0:0:0 | chr7:47825323 |

**Supplementary Table 5:** Severus insertion sequences have high sequence similarity to regions where Manta mate BNDs map.

| Sample | ID | INS length | Max score | Total score | Query cover | E value | Percent identity | Accession | Sequence name |
|---|---|---|---|---|---|---|---|---|---|
| **Pre-treatment tumour** | severus_INS10452 | 544 | 929 | 3822 | 95% | 0.0 | 99.04% | NG_052801.1 | PKD1L1 |
| | severus_INS10474 | 453 | 675 | 822 | 90% | 0.0 | 96.39% | NG_052801.1 | PKD1L1 |
| | severus_INS10486 | 135 | 191 | 191 | 80% | 3e-46 | 98.18% | AC005598.6 | chr4 |
| | severus_INS10502 | 223 | 302 | 302 | 81% | 2e-77 | 96.70% | NG_052801.1 | PKD1L1 |
| | severus_INS10518 | 363 | 499 | 4308 | 80% | 1e-138 | 99.63% | NG_052801.1 | PKD1L1 |
| | severus_INS10559 | 342 | 499 | 499 | 82% | 1e-138 | 98.92% | NG_052801.1 | PKD1L1 |
| | severus_INS10635 | 295 | 520 | 520 | 96% | 7e-145 | 99.65% | NG_052801.1 | PKD1L1 |
| **Post-treatment tumour** | severus_INS10584 | 522 | 436 | 436 | 76% | 1e-117 | 86.99% | NG_052801.1 | PDK1L1 |

**Supplementary Table 6:** Proportions of genomic features present within the classes of differentially modified regions (D(h)MRs).

| Comparison | D(h)MR | Promoters (%) | Exons (%) | Introns (%) | CpG islands (%) | Enhancers (%) | Alu (%) | L1 (%) | Other repeats (%) |
|---|---|---|---|---|---|---|---|---|---|
| **Pre-treatment adj. normal vs Post-treatment adj. normal** | **TOTAL** | **241 (100)** | **520 (100)** | **836 (100)** | **287 (100)** | **624 (100)** | **254 (100)** | **89 (100)** | **567 (100)** |
| | Hypermethylated | 63 (26.7) | 154 (29.6) | 217 (26.0) | 122 (42.5) | 175 (27.3) | 21 (8.3) | 13 (14.6) | 127 (22.4) |
| | Hypomethylated | 164 (69.5) | 356 (68.5) | 605 (72.4) | 162 (56.5) | 458 (71.3) | 225 (88.6) | 73 (82.0) | 428 (75.5) |
| | Hyperhydroxymethylated | 7 (3.0) | 8 (1.5) | 12 (1.4) | 3 (1.0) | 8 (1.3) | 7 (2.8) | 0 (0.0) | 10 (1.8) |
| | Hypohydroxymethylated | 2 (0.9) | 2 (0.4) | 2 (0.2) | 0 (0.0) | 1 (0.2) | 1 (0.4) | 3 (3.4) | 2 (0.4) |
| **Pre-treatment adj. normal vs Pre-treatment tumour** | **TOTAL** | **2031 (100)** | **4170 (100)** | **6185 (100)** | **2644 (100)** | **4313 (100)** | **1767 (100)** | **1147 (100)** | **5062 (100)** |
| | Hypermethylated | 1265 (62.3) | 2535 (60.8) | 2692 (43.5) | 2215 (84.0) | 2020 (46.8) | 208 (11.8) | 78 (6.8) | 2262 (44.7) |
| | Hypomethylated | 764 (37.6) | 1632 (39.1) | 3484 (56.3) | 426 (16.1) | 2292 (53.1) | 1555 (88.0) | 1068 (93.1) | 2795 (55.2) |
| | Hyperhydroxymethylated | 1 (0.1) | 3 (0.1) | 6 (0.1) | 2 (0.1) | 0 (0.0) | 3 (0.2) | 1 (0.1) | 0 (0.0) |
| | Hypohydroxymethylated | 1 (0.1) | 0 (0.0) | 3 (0.1) | 1 (0.0) | 1 (0.0) | 2 (0.1) | 0 (0.0) | 5 (0.10) |
| **Pre-treatment adj.normal vs Post-treatment tumour** | **TOTAL** | **1205 (100)** | **2486 (100)** | **4920 (100)** | **1133 (100)** | **4059 (100)** | **1531 (100)** | **556 (100)** | **3874 (100)** |
| | Hypermethylated | 310 (25.7) | 656 (26.4) | 1126 (22.9) | 411 (36.3) | 731 (18.0) | 338 (22.1) | 100 (18.0) | 731 (18.9) |
| | Hypomethylated | 852 (70.7) | 1734 (69.8) | 3518 (71.5) | 691 (61.0) | 3136 (77.3) | 966 (3.1) | 435 (78.2) | 2993 (77.3) |
| | Hyperhydroxymethylated | 43 (3.6) | 95 (3.8) | 273 (5.6) | 30 (2.7) | 191 (4.7) | 226 (14.8) | 20 (3.6) | 149 (3.9) |
| | Hypohydroxymethylated | 0 (0.0) | 1 (0.0) | 3 (0.1) | 1 (0.1) | 1 (0.0) | 1 (0.1) | 1 (0.2) | 1 (0.0) |
| **Post-treatment adj. norm vs Post-treatment tumour** | **TOTAL** | **1556 (100)** | **3135 (100)** | **5943 (100)** | **1448 (100)** | **4754 (100)** | **2316 (100)** | **714 (100)** | **4980 (100)** |
| | Hypermethylated | 728 (46.8) | 1536 (49.0) | 2731 (46.0) | 677 (46.8) | 1951 (41.0) | 1400 (60.5) | 365 (51.1) | 2458 (49.4) |
| | Hypomethylated | 778 (50.0) | 1516 (48.4) | 2991 (50.3) | 736 (50.8) | 2629 (55.3) | 703 (30.4) | 337 (47.2) | 2431 (48.8) |
| | Hyperhydroxymethylated | 49 (3.2) | 81 (2.6) | 219 (3.7) | 34 (2.4) | 174 (3.7) | 213 (9.2) | 12 (1.7) | 89 (1.8) |
| | Hypohydroxymethylated | 1 (0.1) | 2 (0.1) | 2 (0.0) | 1 (0.1) | 0 (0.0) | 0 (0.0) | 0 (0.0) | 2 (0.0) |
| **Pre-treatment tumour vs Post-treatment tumour** | **TOTAL** | **1465 (100)** | **2816 (100)** | **4368 (100)** | **2027 (100)** | **3052 (100)** | **1193 (100)** | **518 (100)** | **3142 (100)** |
| | Hypermethylated | 241 (16.5) | 525 (18.6) | 1243 (28.5) | 148 (7.4) | 681 (22.3) | 605 (50.7) | 381 (73.6) | 812 (25.8) |
| | Hypomethylated | 1168 (79.3) | 2185 (77.6) | 2785 (63.8) | 1843 (90.9) | 2135 (70.0) | 294 (24.6) | 108 (20.9) | 2154 (68.6) |
| | Hyperhydroxymethylated | 55 (3.8) | 104 (3.7) | 338 (7.7) | 36 (1.8) | 236 (7.7) | 2293 (24.6) | 29 (5.6) | 176 (5.6) |
| | Hypohydroxymethylated | 1 (0.1) | 2 (0.1) | 2 (0.1) | 0 (0.0) | 0 (0.0) | 1 (0.1) | 0 (0.0) | 0 (0.0) |

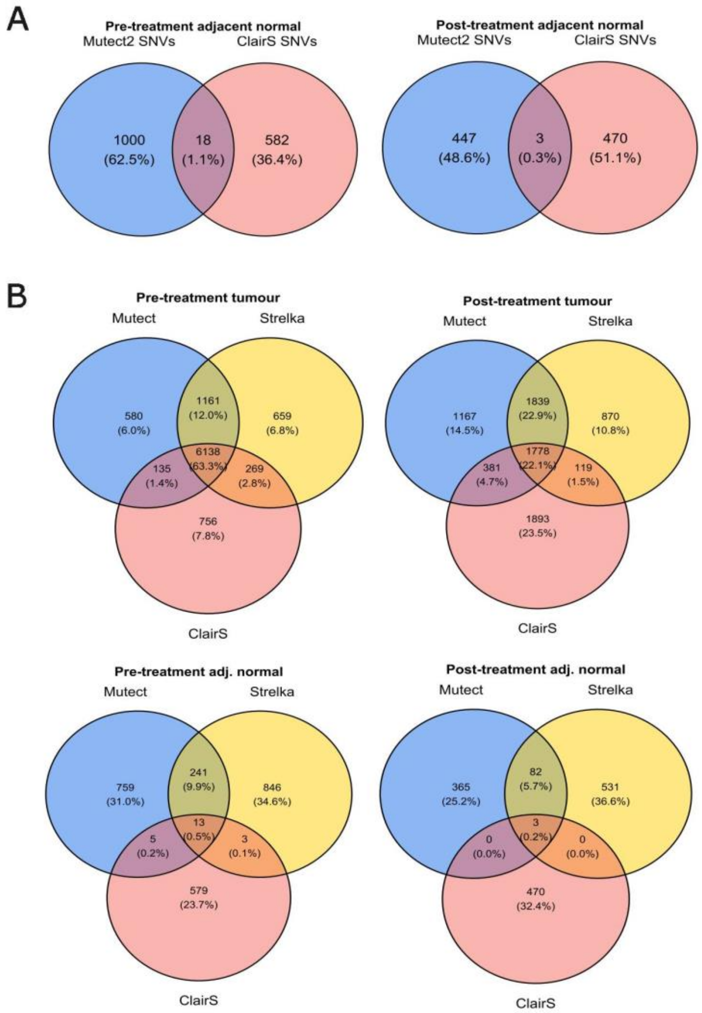


**Supplementary Figure 1. A)** Comparison of SNVs detected by Mutect and ClairS in adjacent normal samples. **B)** Comparison of SNVs detected by Mutect, ClairS and Strelka in tumour and adjacent normal samples

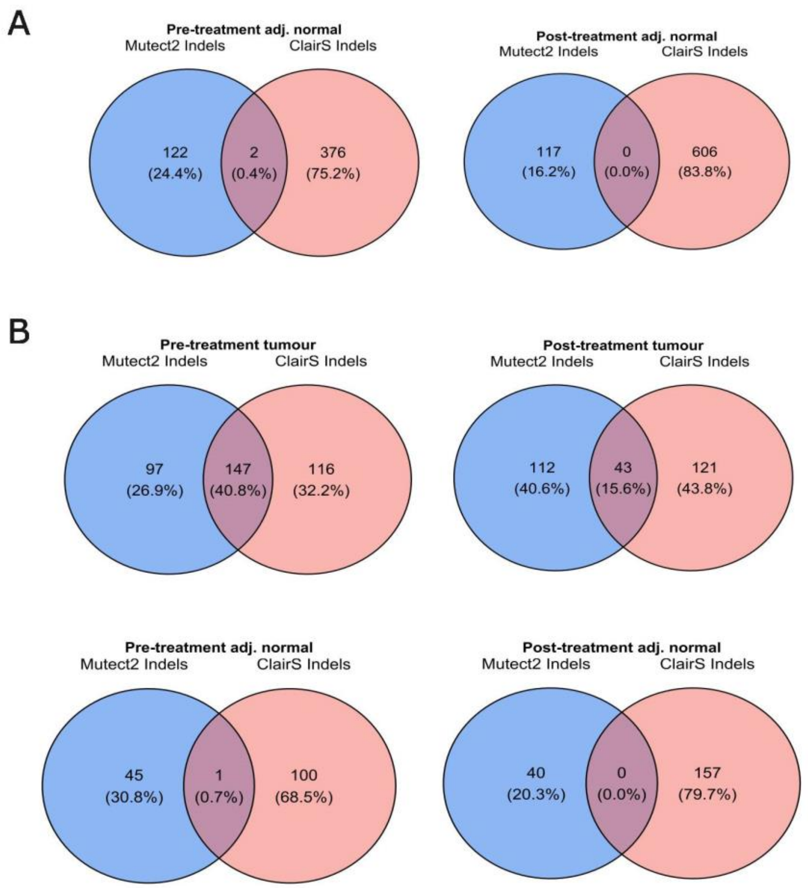


**Supplementary Figure 2. A)** Indel concordance between callers for pre-treatment (left) and post-treatment (right) adjacent normal tissues. **B)** Indel concordance when homopolymers are excluded.

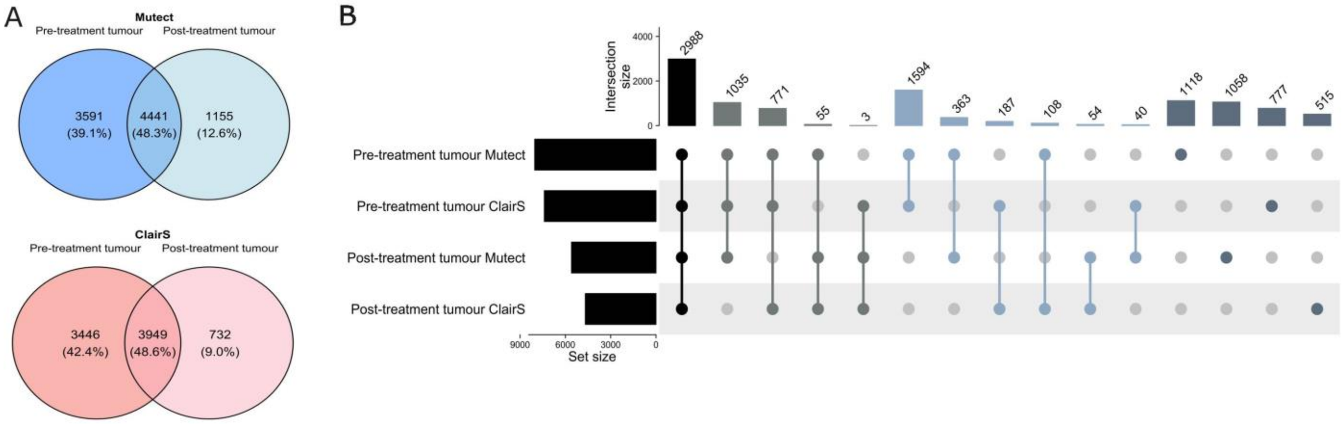


**Supplementary Figure 3. A)** Concordance of SNVs detected by Mutect and ClairS in pre-treatment and post-treatment tumours when low quality SNVs confirmed by orthogonal sequencing data across multiple samples are included. **B)** UpSet plot representing concordance of SNVs detected by Mutect and ClairS across pre- and post-treatment tumours when low quality SNVs confirmed by orthogonal sequencing data across multiple samples are included.

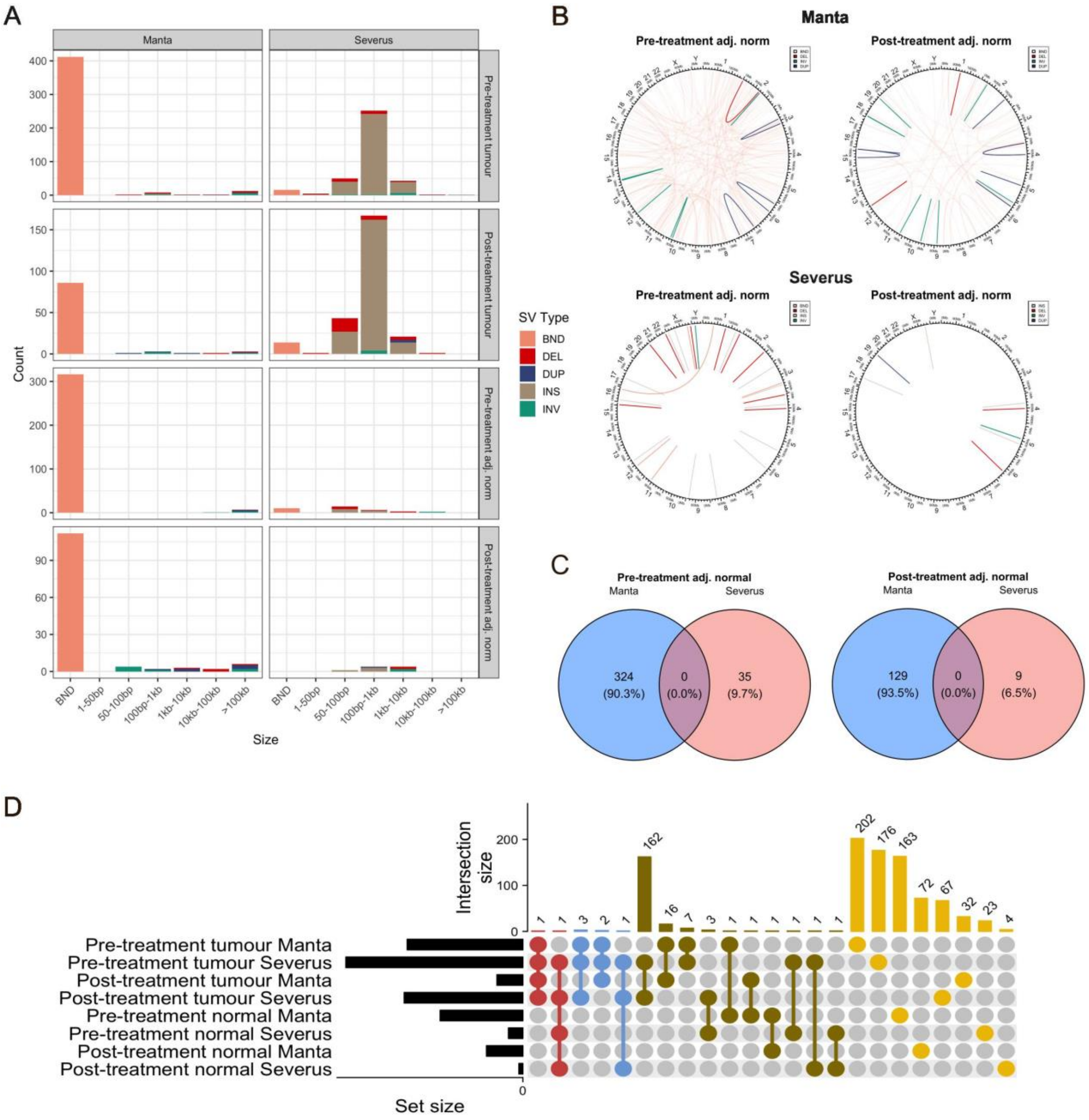


**Supplementary Figure 4. A)** Distribution of SV sizes detected by Manta and Severus by sample. **B)** Strucutral variants detected in the pre- and post-treatment adjacent normal tissues by Manta and Severus. **C)** Minda concordance of SVs between Manta and Severus for the pre- and post-treatment adjacent normal tissues. **D)** Minda concordance of SVs across all samples and SV callers.

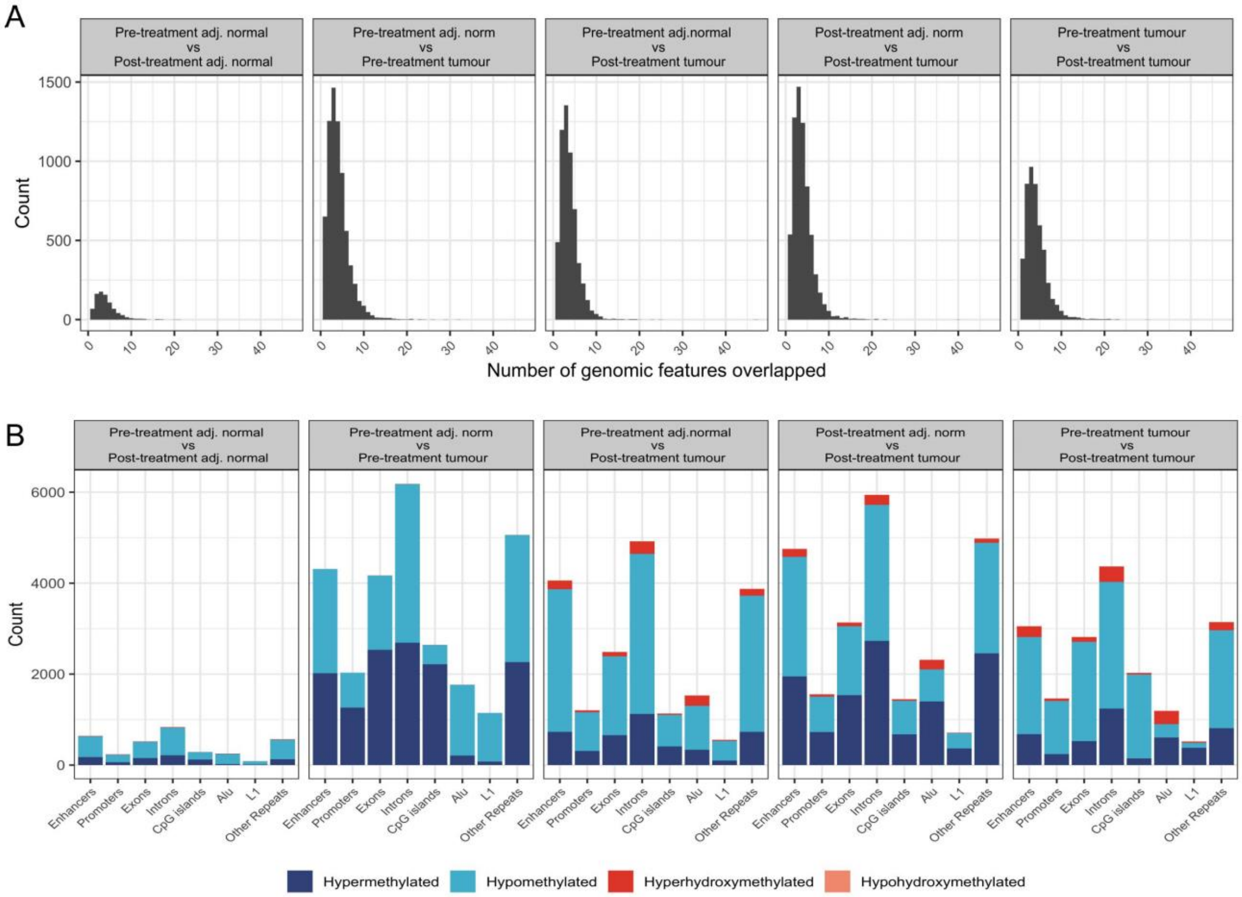


**Supplementary Figure 5. Differentially modified regions overlapped multiple genomic features. A)** Distribution of the number of genomic features within a given DMR. **B)** Breakdown of classes of differentially modified regions that overlap genomic features.
